# An architectural switch in the evolution of the γ-tubulin ring complex

**DOI:** 10.64898/2026.08.03.742184

**Authors:** Rościsław Krutyhołowa, Yi Xie, Banyon Carnell, Hugo Muñoz-Hernández, Daniel Zhang, Florina Marxer, Shaul Yogev, Michal Wieczorek

## Abstract

Canonical microtubules contain 13-protofilaments and are templated by the γ-tubulin ring complex (γ-TuRC). However, some eukaryotes assemble non-canonical microtubules, like the 11-protofilament structures found in *Caenorhabditis elegans*. How γ-TuRCs adapt to template alternative microtubule geometries is unclear. Here, we present the cryo-electron microscopy structure of the *C. elegans* γ-TuRC (γ-TuRC*_Ce_*), revealing a cone-shaped assembly consistent with an 11-protofilament template. While the complex incorporates the conserved subunits actin, GCP2 and GCP3, γ-TuRC*_Ce_* replaces GCP4-6 with a divergent 4-spoked assembly containing additional copies of GCP2 and the nematode-specific proteins GTAP-1 and GTAP-2. Structures of nucleotide-free γ-TuRC*_Ce_* subcomplexes reveal partial γ-tubulin unfolding, suggesting nucleotide binding stabilizes eukaryotic tubulins. Remarkably, reconstituted 4-spoked assemblies can multimerize into ∼13-fold symmetric microtubule nucleation templates *in vitro*, contrasting with the native complex’s 11-protofilament architecture. Our work defines the structural blueprint of an 11-protofilament microtubule template and shows how divergent γ-tubulin components are repurposed to accommodate non-canonical microtubule lattices.

## Introduction

Microtubules are dynamic cytoskeletal polymers that underpin essential cellular processes, including chromosome segregation, intracellular transport, and the establishment of cell shape. Microtubule nucleation is mediated by the γ-tubulin ring complex (γ-TuRC), which provides a pseudo-13-fold symmetric assembly platform that templates the polymerization of microtubules and remains bound at filament minus ends as a stabilizing “cap” (*1–3*). As a result, eukaryotic cells examined to date typically contain 13-protofilament microtubules (*4–6*), yet there are clear examples where this rule is broken (*7*), including in the cortical microtubules of certain parasites (*8*), in the nerve cord microtubules of certain crustaceans (*9*), and in the microtubules of adenosine diphosphate (ADP)-treated platelets (*10*). While α/β-tubulin isotypes have been implicated in regulating microtubule architecture (*11–14*), whether and how the γ-TuRC can accommodate non-canonical microtubule lattices has not been fully investigated.

*Caenorhabditis elegans* offers an attractive model system to study this question, as it has been reported to contain microtubules with at least three different architectures: i) 15 protofilaments in the microtubule bundles of touch receptor neurons (*11*, *15*); ii) 13 protofilaments in the A-microtubules of centrioles (*15*, *16*); and iii) 11 protofilaments in most remaining microtubules (*17*, *18*), making this the canonical lattice architecture in the organism. It is not clear how the 13-protofilament A-microtubules in *C. elegans* centrioles might be specified, but the presence of the 15-protofilament microtubules has been shown to depend on specific α- and β-tubulin isotypes (*11*, *19*). Recently, *in situ* cryo-electron tomography (cryo-ET) coupled with subtomogram averaging (STA) demonstrated that mitotic centrosomes in *C. elegans* embryos contain cone-shaped structures that resemble γ-TuRCs and cap the minus ends of 11-protofilament microtubules emanating from pericentriolar material (*16*). While this study reported an 11-fold rotational symmetry in the capping structure, the limited resolution of the STA reconstruction (∼37 Å) was insufficient to describe the biochemical identity and organization of the proteins in the cap. Thus, whether these caps are truly γ-TuRCs, and if so, how their subunits are organized to nucleate the 11-protofilament architecture of microtubules emanating from *C. elegans* centrosomes, are not known.

In vertebrates, the structure of the γ-TuRC is characterized by three main subassemblies: i) multiple γ-tubulin small complexes (γ-TuSCs) containing γ-tubulin, γ-tubulin complex protein 2 (GCP2), and GCP3; ii) a GCP4/5/4/6 subcomplex, which incorporates γ-tubulin and GCP4-6; and iii) the “lumenal bridge”, where MZT1 and actin associate with the N-terminal domains of GCP3 (GCP3-NTD) and GCP6 (GCP6-NTD) (*20–24*). Central to the γ-TuRC’s function is γ-tubulin, which binds and hydrolyzes GTP much like its structural homolog β-tubulin does within the microtubule lattice (*25–27*). Within the complex, each γ-tubulin is bound to a conserved C-terminal GCP GRIP2 domain to form a “spoke”, and 14 of these γ-tubulin:GCP spokes associate laterally via the N-terminal GCP GRIP1 domains to form the γ-TuRC. Five γ-TuSCs are located at spoke positions 1-2, 3-4, 5-6, 7-8, and 13-14 in the cone-shaped structure, while a GCP4/5/4/6 subassembly occupies positions 9-12. Because spoke 14 partially overlaps with spoke 1, exactly 13 γ-tubulin subunits remain available to bind α-tubulin. The asymmetric organization of GCPs in the vertebrate γ-TuRC is thought to be coordinated in part by the lumenal bridge (*21*, *28*). Thus far, however, only vertebrate γ-TuRCs have been shown to build an actin-containing lumenal bridge; whether actin is also a component of the *C. elegans* γ-TuRC is unclear. More broadly, how the composition, stoichiometry, and relative organization of subunits in the *C. elegans* γ-TuRC may have been modified to accommodate non-canonical microtubules is an open question.

Previous work has identified the *C. elegans* homologs of the γ-TuSC proteins γ-tubulin (TBG-1), GCP2 (GIP-2), and GCP3 (GIP-1) (*29*). *C. elegans* also contains a homolog of MZT1 (MZT-1), which has been shown to associate with GCP3 (*30*, *31*). Similarly, SPD-5, an ortholog of *D. melanogaster* centrosomin (Cnn), human CDK5RAP2, and budding yeast Spc110p, is a centrosomal protein potentially important for γ-TuRC assembly and activity in *C. elegans* that has been reported to associate with γ-TuSCs in a phosphorylation-dependent manner (*32*, *33*). Although homologs of GCP4, GCP5 or GCP6 proteins in *C. elegans* have not been identified, a recent study characterized two genes whose protein products are physically and functionally associated with γ-tubulin, termed γ-tubulin associated protein 1 (GTAP-1) and GTAP-2 (*31*). These proteins contain the conserved GRIP1 and GRIP2 domains shared by other GCP members, but their NTDs do not share obvious homology with MZT1-binding domains found in GCP5 and GCP6. Moreover, only two such GCP homologs have been identified thus far, yet *C. elegans* lacks three of the GCPs found in vertebrates (GCP4, 5 and 6). Thus, the role of the GTAP-1/2 proteins in the assembly and architecture of *C. elegans* γ-tubulin complexes remains unclear.

Here, we use cryo-EM and biochemical reconstitutions to investigate the structure, composition, and assembly of *C. elegans* γ-tubulin complexes. We show that the native *C. elegans* γ-TuRC is built from γ-tubulin, GCP2, GCP3, GTAP-1, GTAP-2, and actin, which together form a cone-shaped structure consistent with an ∼11-protofilament microtubule nucleation template. A reconstituted subcomplex consisting of γ-tubulin, GCP2, GTAP-1, and GTAP-2 resembles the GCP4/5/4/6 assembly found in vertebrates, indicating that *C. elegans* GCP2 has replaced the role of vertebrate GCP4 in this organism. Remarkably, these subcomplexes can multimerize into γ-TuRC-like structures that can nucleate microtubules but now adopt a ∼13-protofilament template architecture. Our results not only show how the *C. elegans* γ-TuRC has been evolutionarily repurposed to accommodate non-canonical 11-protofilament microtubules, but also reveal that γ-tubulin subcomplexes from the same species can form microtubule nucleation templates with either ∼11- or ∼13-protofilament-like geometries.

## Results

### The C. elegans γ-TuRC adopts an ∼11-fold symmetric architecture and is assembled from an atypical set of components

To elucidate the structure of the *C. elegans* γ-TuRC (hereafter, γ-TuRC*_Ce_*), we isolated the native complex from a CRISPR/Cas9-edited *C. elegans* strain expressing the γ-tubulin ortholog TBG-1 (hereafter, γ-tubulin*_Ce_*) fused to a C-terminal FLAG tag (Figure 1A). Tagging did not affect animal viability or fertility, suggesting that it does not interfere with γ-TuRC*_Ce_*function (*31*). Mass spectrometry (MS) of γ-TuRC*_Ce_* isolated from whole animal lysates confirmed the presence of γ-tubulin*_Ce_*, the GCP2 ortholog GIP-2 (hereafter, GCP2*_Ce_*), the GCP3 ortholog GIP-1 (hereafter, GCP3*_Ce_*), as well as the nematode-specific <u>γ-t</u>ubulin*_Ce_*-<u>a</u>ssociated <u>p</u>rotein (GTAP) 1 and GTAP-2 (Supplementary Table 1) (*31*). Although the *C. elegans* orthologs of MZT1 (MZT1*_Ce_*) and CDK5RAP2/Cnn (SPD-5) have previously been reported to interact with *C. elegans* γ-tubulin complexes (*30–32*), we did not identify these proteins by MS, which may be due to the small size of MZT1*_Ce_* (∼7 kDa), the reported dispensability of MZT1*_Ce_* for the function of certain γ-tubulin complexes (*30*, *31*), or differences between complexes obtained from whole animals (our study) vs. from embryos (*31*). Total internal reflection fluorescence (TIRF) microscopy showed that immobilized γ-TuRC*_Ce_* nucleated microtubules in the presence of fluorescently-labeled porcine brain tubulin and 1 mM GTP (Supplementary Figure 1A-B). Microtubules grew from only one end and exhibited polymerization rates consistent with plus (fast) rather than minus (slow) end growth (*34*). This suggests that γ-TuRC*_Ce_* templates the assembly of filament plus ends and remains bound to their minus ends as a stabilizing cap, consistent with TIRF microscopy of vertebrate γ-TuRCs from other species (*23*, *35–37*).

**Figure 1.**
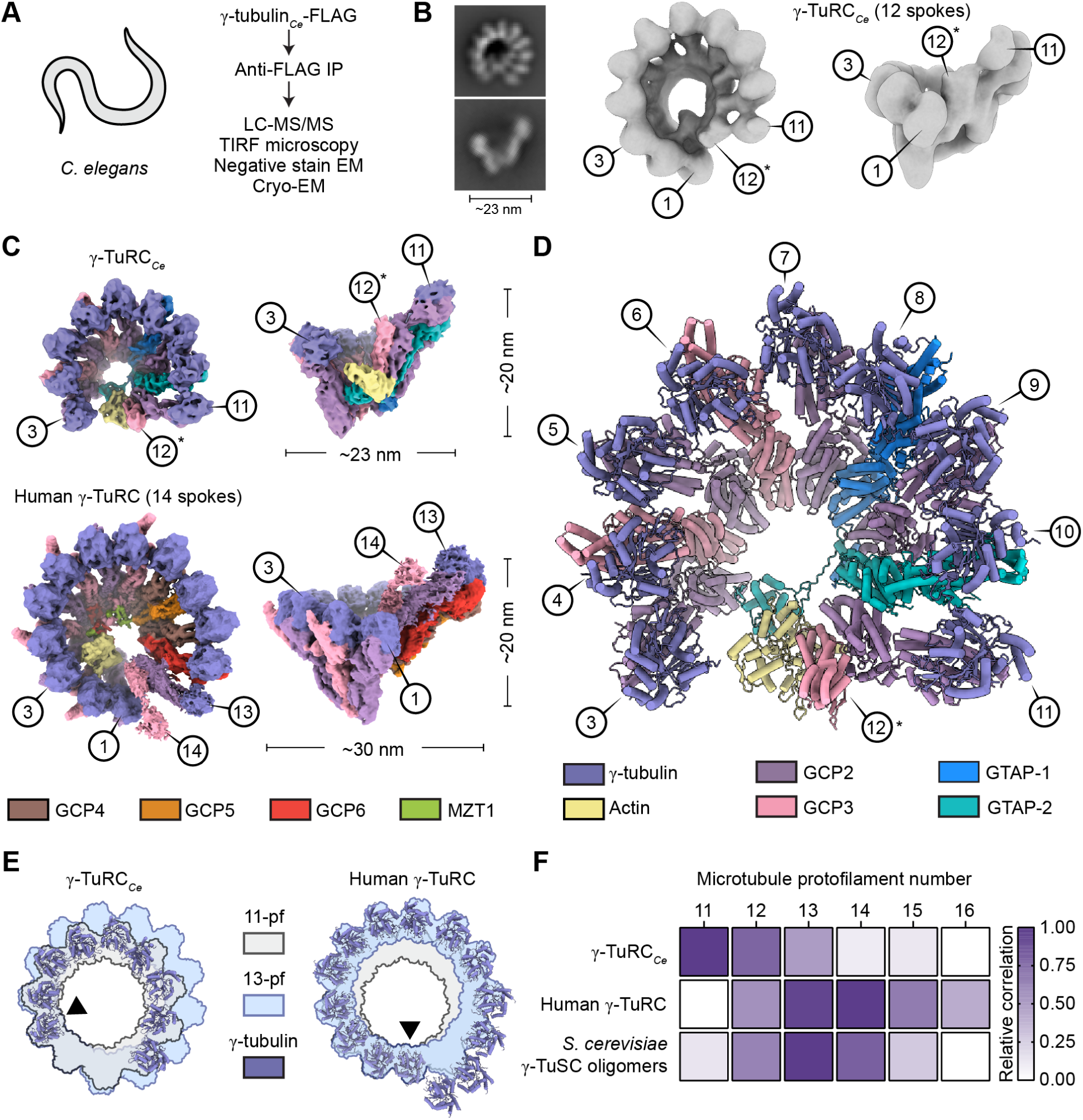
γ-TuRC*_Ce_* adopts an ∼11-fold symmetric architecture and is assembled from an atypical set of components. **A)** Workflow for γ-TuRC*_Ce_* isolation and characterization from *C. elegans* whole animals. **B)** Left: γ-TuRC*_Ce_* negative stain EM 2D class averages. The diameter of the complex as estimated from the 2D averages is indicated. Right: Top and side views of the γ-TuRC*_Ce_* negative stain EM 3D reconstruction. γ-TuRC*_Ce_* spokes 1, 3, 11 and 12 are indicated. **C)** Top and side views of γ-TuRC*_Ce_* (top; this study) and human γ-TuRC (bottom; EMD-21073 low-pass filtered to 6.5 Å (*20*)) cryo-EM density maps. Estimates of complex widths and heights are indicated. γ-TuRC*_Ce_* subunits are colored according to the legend at the bottom of panels **C)** to **D)**. Spokes 1, 3, 11, 12, 13 and/or 14 are indicated for both complexes, where applicable. **D)** Cartoon representation of γ-TuRC*_Ce_*built using the cryo-EM reconstruction in **C)**. Spoke numbers are indicated and subunits are colored according to the legend. γ-tubulin, GCP2 and GCP3 in the legend also refer to the *C. elegans* homologs. Asterisks in panels **B) - D)** indicate the partially-resolved spoke at γ-TuRC*_Ce_* position 12. **E)** ã-tubulin rings from γ-TuRC*_Ce_* (left) and human γ-TuRC (right; PDB ID: 6V6S (*20*)) models aligned to a ring of β-tubulin from an 11-protofilament microtubule (grey; EMD-5191) or a 13-protofilament microtubule (light blue; EMD-5193) (*38*). The γ- and β-tubulin subunits used for alignments are indicated with a black triangle. **F)** Relative cross-correlation values of β-tubulin rings from various microtubule protofilament number architectures fitted to γ-tubulin rings from γ-TuRC*_Ce_* (this study), as well as human γ-TuRC in the “open” conformation (PDB ID: 6V6S (*20*)), and budding yeast γ-TuSC oligomers in the “closed” conformation (PDB ID: 5FLZ (*39*)) as controls, colored according to the legend on the right. For visualization purposes, values are normalized such that the maximum in each comparison corresponds to 1, while minimum values correspond to 0 (see Methods).

Negative stain EM of γ-TuRC*_Ce_* revealed particles with a “lock washer” geometry resembling γ-TuRCs from other species (Supplementary Figure 1C) (*22*, *35*). Single particle analysis (SPA) of the negative stain EM data yielded a 3D reconstruction of γ-TuRC*_Ce_* at ∼19 Å resolution (FSC_0.143_; Figure 1B and Supplementary Figure 1D-E). The reconstruction shows that γ-TuRC*_Ce_* adopts a 12-spoked, cone-shaped assembly measuring ∼23 nm in diameter (Figure 1B), which is smaller than γ-TuRCs from other species (e.g.,14 spokes and ∼30 nm diameter for human γ-TuRC; Figure 1C) (*20*, *22*). Notably, density for the top half of the 12th spoke in γ-TuRC*_Ce_* could not be resolved by negative stain EM (Figure 1B and Supplementary Figure 1D-E). To assess the biochemical composition of γ-TuRC*_Ce_*, we employed SPA of vitrified complexes imaged by cryo-EM, which yielded a 3D reconstruction at 6.1 Å resolution (FSC_0.143_; Figure 1C, Supplementary Table 2 and Supplementary Figure 2A-D). The cryo-EM reconstruction is qualitatively consistent with the 12-spoked reconstruction determined by negative stain EM (Figure 1B), except that density for spokes 1-2 is missing. These differences are likely explained by the previously reported stabilizing effects of uranyl salts on protein complexes (*40*), and suggest that γ-TuRC*_Ce_* may be less stable compared to native γ-TuRCs from vertebrates (*20*, *22*, *23*).

Systematic fitting of AlphaFold-predicted models of γ-TuRC*_Ce_* subunits into the cryo-EM density map revealed the organization of the *C. elegans*-specific GCP-containing spokes (Supplementary Figure 3A) (*41*), allowing us to build a molecular model of the complex (Figure 1D, Supplementary Figure 3B and Supplementary Table 3). The cryo-EM-derived γ-TuRC*_Ce_* model shows that spokes 3-6 consist of a pair of γ-TuSCs containing γ-tubulin*_Ce_*, GCP2*_Ce_* and GCP3*_Ce_*, arranged in a side-by-side fashion similar to γ-TuRCs in other species (*20*, *22–24*, *42*, *43*). Unlike other γ-TuRCs, however, these subunits are followed by an arrangement of γ-tubulin*_Ce_*-associated GCP2*_Ce_*/GTAP-1/GCP2*_Ce_*/GTAP-2 occupying spokes 7-10 of γ-TuRC*_Ce_*. This “4-spoked assembly” is then followed by a final γ-TuSC at spokes 11-12 to complete γ-TuRC*_Ce_*. Consistent with the negative stain EM reconstruction (Figure 1B and Supplementary Figure 3C), γ-tubulin*_Ce_* and the associated GRIP2 domain of the GCP subunit in the final, 12th spoke of the complex are also poorly resolved by cryo-EM (Figure 1C-D), suggesting substantial flexibility at this particular GCP subunit. Despite this flexibility, the map-to-model cross-correlation of the well-resolved GRIP1 domain at this position in the cryo-EM map supports its assignment as GCP3*_Ce_* (Supplementary Figure 3A-B).

Next, to assess which microtubule architecture γ-TuRC*_Ce_* is most likely to specify, we performed two different analyses of the complex, with similar outcomes: i) cross-correlation of the γ-tubulin*_Ce_* “ring” with published structures of microtubules exhibiting a variety of protofilament numbers (*38*); and ii) rotational averaging of the entire complex (see Methods). The results indicate that the structure of γ-TuRC*_Ce_* matches best with the 11-protofilament lattice of *C. elegans* cytoplasmic microtubules (Figure 1E-F and Supplementary Figure 3D-E). Moreover, the γ-TuRC*_Ce_* model is qualitatively consistent with a recent ∼37 Å resolution STA reconstruction of centrosome-embedded microtubules in *C. elegans* embryos (*16*), whose ends are capped by γ-TuRC-like structures displaying 11-fold rotational symmetry (Supplementary Figure 3F). These findings demonstrate that γ-TuRC*_Ce_*adopts an ∼11-fold symmetric architecture and is assembled from an atypical set of GCP subunits.

### The N-terminus of GTAP-2 interacts with C. elegans actin

Analogous to actin in the lumenal bridge of vertebrate γ-TuRCs (*20*, *22*), our cryo-EM reconstruction also revealed the presence of an actin molecule in the lumen of γ-TuRC*_Ce_* (actin*_Ce_*; Figure 1C-D and Supplementary Figure 4A-B). In vertebrate γ-TuRCs, the lumenal bridge is composed of MZT1 in complex with the <u>N-T</u>erminal <u>D</u>omain of GCP3 (MZT1:GCP3-NTD), as well as MZT1:GCP6-NTD, which binds directly to actin and positions it ∼29 Å away from the GRIP1 domain of the terminal GCP3 subunit (Figure 2A-B). In γ-TuRC*_Ce_*, however, actin*_Ce_*is rotated by ∼13° compared to its human counterpart, placing it in direct contact with the GRIP1 domain of GCP3*_Ce_* at position 12 (Figure 2A-B). The barbed end groove of actin*_Ce_* associates with the N-terminal domain of GTAP-2 (GTAP-2-NTD) (Supplementary Figure 4B-C), which can be traced directly to GTAP-2’s GRIP1 domain (Figure 1D). Notably, density for MZT1*_Ce_* was not observed in any part of our cryo-EM reconstruction. Consistent with this observation, an AlphaFold3 prediction supports a direct interaction between GTAP-2-NTD and actin*_Ce_* in a MZT1-independent manner (Supplementary Figure 4A), with the actin-binding region of GTAP-2-NTD and MZT1:GCP6-NTD sharing similar secondary structure features (Supplementary Figure 4C).

**Figure 2.**
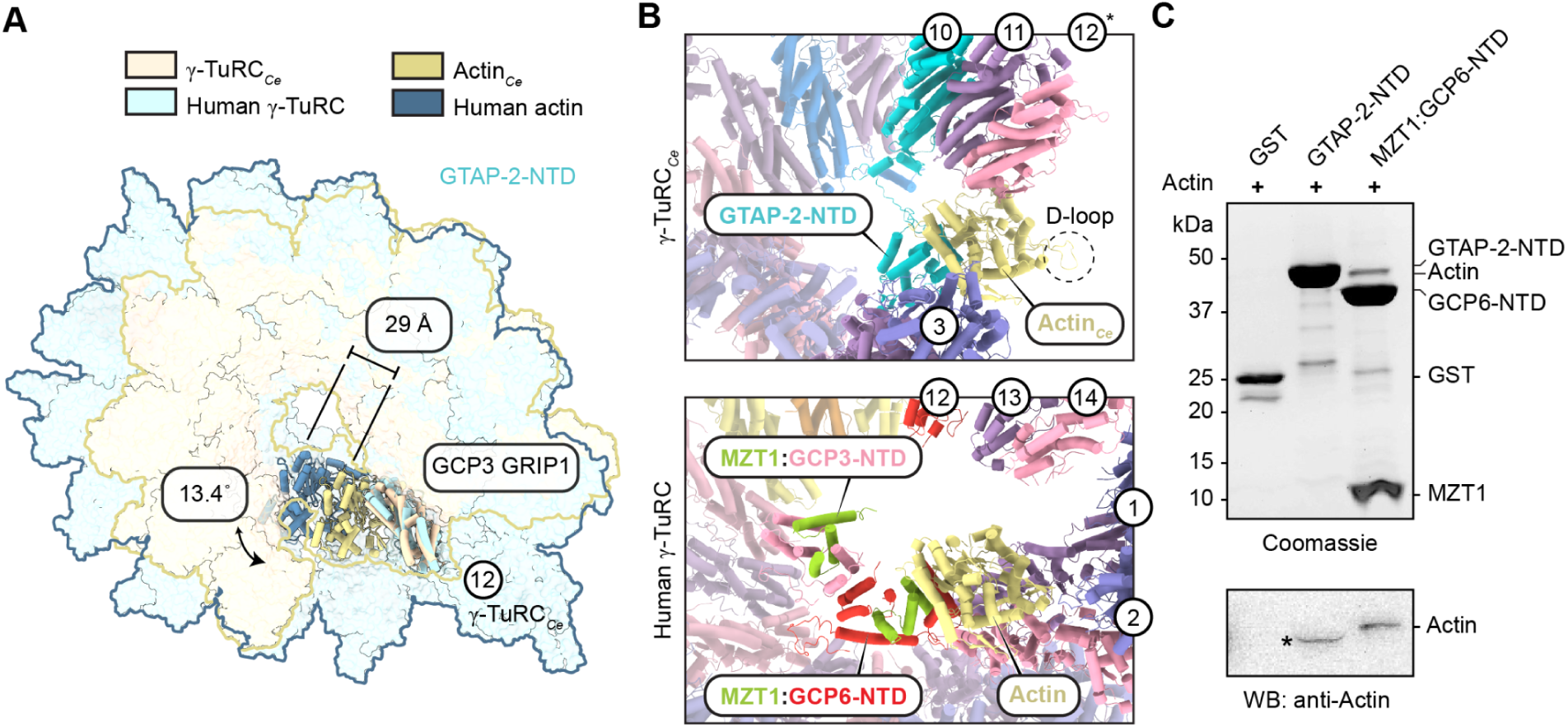
The N-terminus of GTAP-2 interacts with *C. elegans* actin. **A)** Top view of the γ-TuRC*_Ce_* model aligned to the human γ-TuRC model using the GCP3 GRIP1 domains of the last spoke of each complex. Relative displacement and rotation of actin between the two complexes is indicated, with actin and the aligned GCP3 GRIP1 domains shown in cartoon representation, while the rest of the complex is shown in transparent surface representation. Complexes and subunits are colored according to the legend at the top. **B)** Zoomed-in and rotated views of actin in γ-TuRC*_Ce_* (top) and in human γ-TuRC*_Hs_* (bottom). Models were aligned to one another via actin. Lumenal bridge components, actin-interacting γ-TuRC spoke numbers, and position numbers of the last three spokes of each complex are indicated. All other subunits are rendered with a transparency of 50% for figure clarity. Subunits are colored according to the legend in Figure 1C-D. The location of actin*_Ce_*’s D-loop is indicated by a dashed circle. **C)** Coomassie-stained SDS-PAGE gel (top) and anti-actin western blot (bottom) of GST pull-down experiments performed with actin and either GTAP-2-NTD or MZT1:GCP6-NTD. Band locations of relevant proteins are indicated. The actin band in the GTAP-2-NTD lane (indicated by an asterisk) is slightly lower than expected because it presumably overlaps with GST-tagged GTAP-2-NTD, which runs at a similar molecular weight as actin.

To further test whether GTAP-2-NTD alone is sufficient for actin binding, we performed pull-down assays using purified proteins fused to glutathione S-transferase (GST), with a GST-tagged human MZT1:GCP6-NTD subcomplex serving as a positive control (*21*). Consistent with the structure of γ-TuRC*_Ce_*, we found that GTAP-2-NTD could pull down unpolymerized mammalian actin under similar conditions as the MZT1:GCP6-NTD control (Figure 2C and Supplementary Figure 4D-E). Together, our findings reveal the presence of a lumenal bridge-like structure in γ-TuRC*_Ce_*, in which *C. elegans* GTAP-2-NTD alone structurally substitutes for the vertebrate MZT1:GCP6-NTD module.

### A 4-spoked assembly containing GCP2_Ce_/GTAP-1/GCP2_Ce_/GTAP-2 is structurally analogous to GCP4/GCP5/GCP4/GCP6 in vertebrates

Given GTAP-2’s obvious structural role in γ-TuRC*_Ce_*, we next focused on the 4-spoked assembly consisting of GCP2*_Ce_*, GTAP-1 and GTAP-2 at positions 7-10 of the complex (Figure 1C-D). We performed a Hidden Markov Model (HMM)-based orthology analysis of γ-TuRC*_Ce_* components compared to closely related eukaryotes (see Methods). The resulting phylogenetic tree suggests that GTAP-1 and GTAP-2 emerged simultaneously with the loss of GCP4-6 in a common ancestor of *C. elegans* and its related species (Supplementary Figure 5A-D).

Subunit-specific searches suggest that the ancestor of GTAP-2 is most likely GCP6-like, consistent with the interaction of GTAP-2-NTD with actin*_Ce_*, and that the ancestor of GTAP-1 is most likely GCP5-like (Supplementary Figure 5E), providing a potential evolutionary origin of these subunits in *C. elegans* (*31*). The loss of GCP4 can be further explained by the structure of γ-TuRC*_Ce_*, which indicates that GCP2*_Ce_* is both GCP2-like (as part of the γ-TuSCs) and GCP4-like (as part of a GCP4/GCP5/GCP4/GCP6-like, 4-spoked subassembly; Figure 3A).

**Figure 3.**
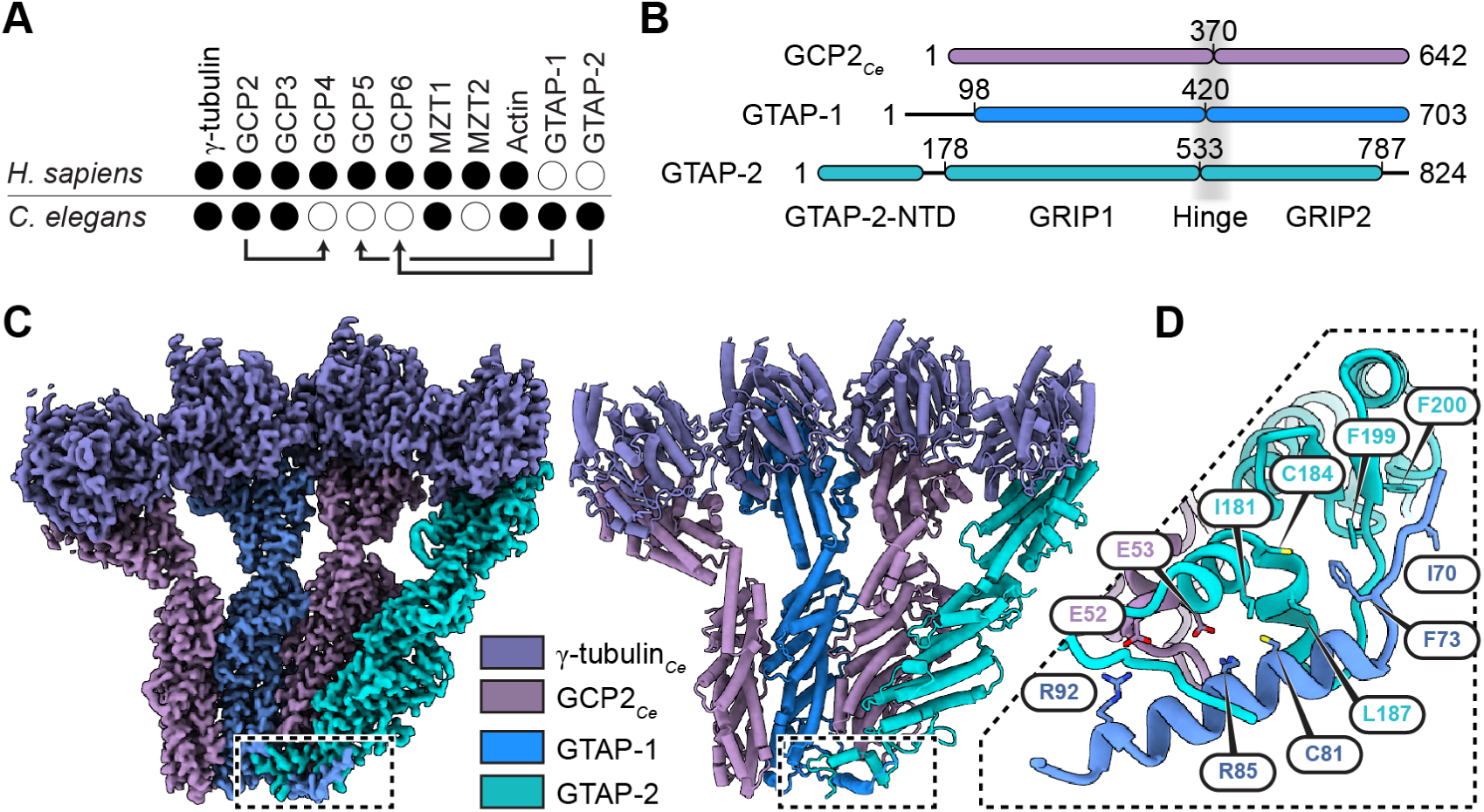
A 4-spoked assembly containing GCP2*_Ce_*/GTAP-1/GCP2*_Ce_*/GTAP-2 is structurally analogous to GCP4/GCP5/GCP4/GCP6 in vertebrate γ-TuRCs. **A)** Schematic proposing evolutionary replacement of *H. sapiens* γ-TuRC subunits with *C. elegans* orthologs, based on bioinformatics and structural analysis of γ-TuRC*_Ce_*. Filled circles indicate the presence of a particular γ-TuRC subunit in the given species. Arrows indicate *C. elegans* subunits (start of arrows) that have functionally replaced corresponding γ-TuRC subunits in *H. sapiens* and other γ-TuRCs studied to date (end of arrows). **B)** Schematic diagrams depicting γ-TuRC*_Ce_* GCP sequences in the reconstituted 4-spoked assembly. GCP domains and features are indicated. **C)** Left: Composite cryo-EM map from multi-body refinement of the 4-spoked assembly colored according to the legend. Maps were postprocessed using EMReady prior to combining (*44*). Right: Cartoon representation of the recombinant 4-spoked assembly model built using the density map on the left. Intertwined segments corresponding to segments of GTAP-1 and GTAP-2, which are N-terminal to the respective GRIP1 domains, are indicated by the dashed box. **D)** Zoomed-in view of the dashed region in **C)**. Relevant residue-residue contacts stabilizing the N-terminal region of GTAP-1 that reaches across GCP2*_Ce_*to interface with GTAP-2 are indicated and shown as stick representation with heteroatom coloring.

To better understand the nature of these divergent GCPs in γ-TuRC*_Ce_*, and to overcome the limited resolution of the EM reconstructions of the native complex, we next sought to reconstitute the *C. elegans* complex in insect cells (Supplementary Figure 6A). Despite our attempts to co-overexpress all candidate γ-TuRC*_Ce_* proteins including GCP3*_Ce_*, MZT1*_Ce_*, and a fragment of the accessory protein SPD-5 homologous to budding yeast Spc110_1-220_ (*45*), only complexes containing full length γ-tubulin*_Ce_*, GCP2*_Ce_*, GTAP-1 and GTAP-2 could be successfully purified (Supplementary Figure 6B). Negative stain EM 2D class averages indicated that these complexes form 4-spoked assemblies resembling the one occupying positions 7-10 in γ-TuRC*_Ce_* (Supplementary Figure 6C). We therefore used cryo-EM to generate a 3D reconstruction at 3.0 Å resolution (FSC_0.143_; Figure 3C, Supplementary Figure 6D-F, and Supplementary Table 4), which confirmed that this subcomplex indeed corresponds to GCP2*_Ce_*/GTAP-1/GCP2*_Ce_*/GTAP-2-containing 4-spoked assemblies (Supplementary Table 5).

The 4-spoked assembly reconstruction revealed several notable features in this evolutionarily divergent γ-tubulin subcomplex. First, an N-terminal portion of GTAP-1 adjacent to the GRIP1 domain (residues ∼69-100) snakes along the bottom of the neighboring GCP2*_Ce_* and partially up along the side of GTAP-2’s GRIP1 domain (Figure 3D). This extended interface comprises an acidic patch on GCP2 (E52 and E53) that complements basic residues in GTAP-1 (R92 and R85), as well as potential hydrophobic interactions between a stretch of residues in GTAP-1 (C81, F73 and I70) and GTAP-2 (I181, C184, L187, F199 and F200). These structural features are absent in the other γ-TuRC*_Ce_* subunit interfaces, suggesting they may contribute to the specific subunit ordering of the 4-spoked assembly in γ-TuRC*_Ce_*.

Second, and in contrast with γ-TuRC*_Ce_*, neither endogenous insect actin, which can co-purify with vertebrate γ-TuRCs (*24*), nor GTAP-2-NTD, are resolved in the 4-spoked assembly reconstruction. This is presumably due to a lack of detectable GCP3*_Ce_*in the purified subcomplexes (Supplementary Figure 6B), highlighting the requirement of GCP3*_Ce_* at position 12 of γ-TuRC*_Ce_* for the actin-containing structural bridge in the native assembly (Figures 1C-D and 2B). Third, the conformation of the reconstituted 4-spoked assembly is nearly identical to that of the 4-spoked assembly found in γ-TuRC*_Ce_* (overall RMSD = 1.0 Å; Supplementary Figure 6G). This correlates with the apparent biochemical stability of the 4-spoked assembly and suggests that GCP2*_Ce_*/GTAP-1/GCP2*_Ce_*/GTAP-2 may constitute part of an initial γ-TuRC*_Ce_* assembly scaffold, as proposed for the GCP4/GCP5/GCP4/GCP6 subcomplex in vertebrate γ-TuRCs (*28*).

### Structures of apo and GTP-bound γ-tubulin_Ce_ reveal a role for nucleotide binding in eukaryotic tubulins

Intriguingly, we did not observe any density for guanine nucleotide in γ-tubulin*_Ce_* subunits in the 4-spoked assembly cryo-EM reconstruction (Figure 4A), even though GTP was present throughout the purification. To our knowledge, a nucleotide-free (“apo”) structure of any eukaryotic tubulin family member has not yet been reported. We therefore compared the model of apo-γ-tubulin*_Ce_*in the 4-spoked assembly with available α-, β- or γ-tubulin structures to investigate how nucleotide binding might influence the overall fold of eukaryotic tubulins (*27*, *46*). The most striking differences in apo-γ-tubulin*_Ce_* vs. nucleotide-bound α-, β- or γ-tubulin are: i) helix H3’ is resolved but unfolded; ii) helix H2 is entirely unresolved; and iii) helices H1 and H5 are partially unresolved, particularly near the active site (Supplementary Figure 7A-C, E). Notably, these secondary structure elements are well-resolved in models of α-, β- and γ-tubulin irrespective of the nucleotide or mimic bound (*27*, *46*).

**Figure 4.**
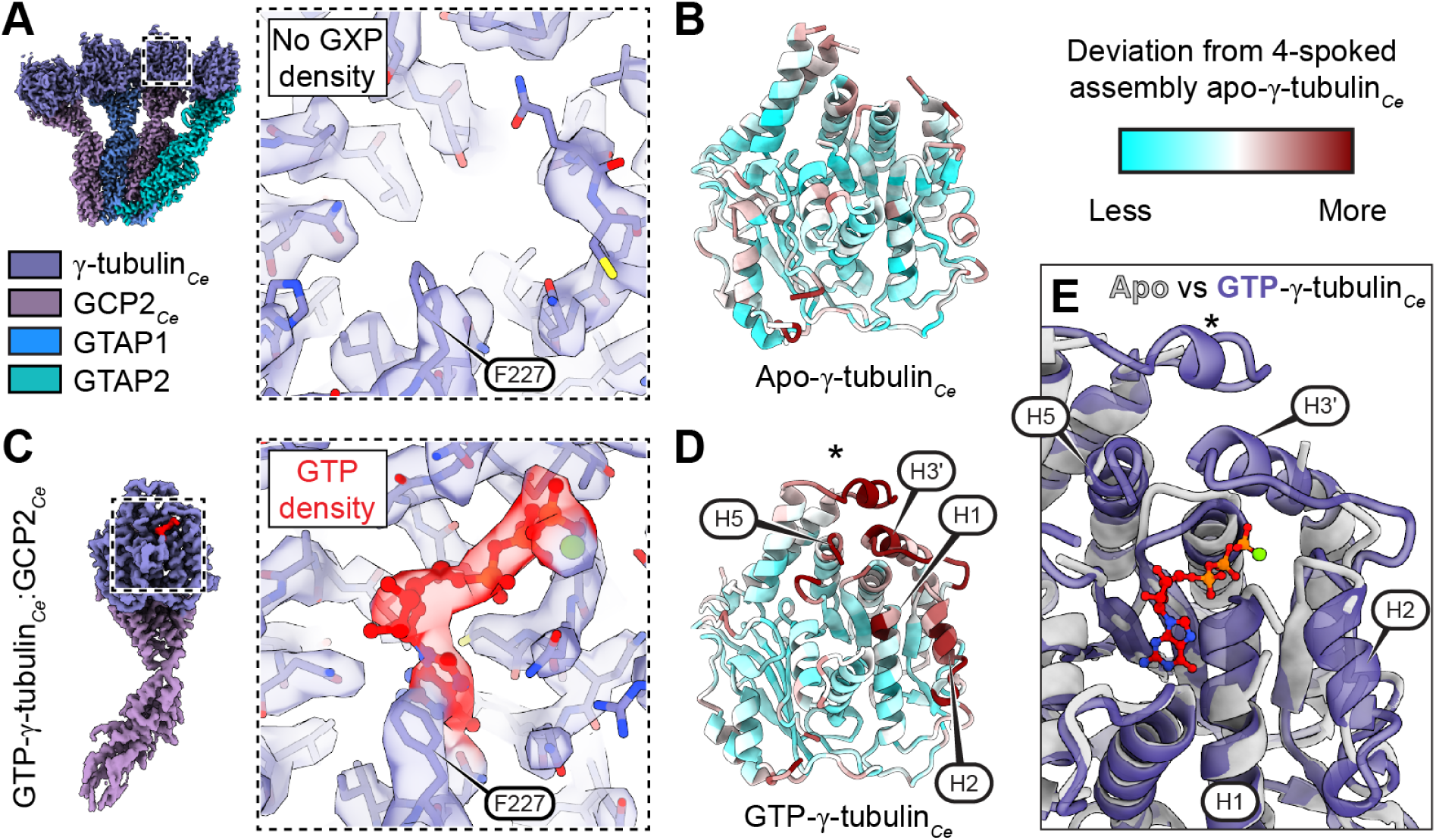
Apo and GTP-bound γ-tubulin*_Ce_* models reveal nucleotide-induced structural rearrangements in eukaryotic tubulins. **A)** Left: Top view of the 4-spoked assembly consensus cryo-EM reconstruction in Figure 3C with a dashed region around the nucleotide binding pocket of a single γ-tubulin*_Ce_* monomer. Right: Zoomed-in view of the γ-tubulin*_Ce_* model (stick representation, colored by heteroatom) in the corresponding cryo-EM density (transparent surface) from the dashed region on the left. **B)** Cartoon representation of apo-γ-tubulin*_Ce_* built using the reconstituted γ-tubulin*_Ce_*:GCP2*_Ce_* density map (Supplementary Figure 8C,F). The model is colored according to the output of the “measure mapvalues” command in ChimeraX, as indicated with the legend on the right and using the 4-spoked assembly density as the reference map. **c)** Left: DynaMight cryo-EM reconstruction of reconstituted γ-tubulin*_Ce_*:GCP2*_Ce_* supplemented with GTP and colored according to the legend in panel **A)** (*47*). The density for guanine nucleotide is colored red for clarity, and a dashed box outlines the nucleotide binding pocket of γ-tubulin*_Ce_*. The map was post-processed using EMReady (*44*). Right: Zoomed-in view of the GTP-γ-tubulin*_Ce_* model (stick representation, colored by heteroatom) in the corresponding cryo-EM density (transparent surface) from the dashed region on the left. The GTP model (stick and ball representation) and the corresponding density are colored in red. A magnesium ion is modeled and shown as a green sphere. Residue F227 is indicated in panels **A)** and **C)** to emphasize that the views of the nucleotide pocket are the same. **D)** Cartoon representation of GTP-γ-tubulin*_Ce_* built using the GTP-supplemented reconstituted γ-tubulin*_Ce_*:GCP2*_Ce_*density map in panel **C)**. The model is colored according to the output of the “measure mapvalues” command in ChimeraX, as in panel **B)**. Locations of helices H1, H2, H3’, and H5 are indicated. **E)** Zoomed-in view of the nucleotide binding pocket comparing apo-γ-tubulin*_Ce_* (from the 4-spoked assembly, grey cartoon representation) with GTP-γ-tubulin*_Ce_* (from panels **C)-D)**, purple transparent cartoon representation). Locations of helices H1, H2, H3’, and H5 are indicated. Asterisks in panels **D)** and **E)** indicate the location of helix H11’, which can be difficult to model even in nucleotide-bound γ-tubulin and was therefore not considered in our analysis (*20*).

The absence of nucleotide in γ-tubulin*_Ce_* could be a distinct property of the 4-spoked assembly, and although the nucleotide pockets also appear empty in the γ-TuRC*_Ce_* reconstruction, the limited resolution of γ-tubulin*_Ce_*densities there did not permit a definitive assessment of γ-tubulin*_Ce_*’s basal nucleotide state. To examine whether the apo state inherent to γ-tubulin*_Ce_*, we reconstituted and purified a single-spoked γ-tubulin*_Ce_*:GCP2*_Ce_* subcomplex from insect cells (Supplementary Figure 8A-B). We used cryo-EM to generate a 3D reconstruction of the ∼125 kDa subcomplex at 5.9 Å resolution from ∼53,000 particles (FSC_0.143_; Figure 4B, Supplementary Figure 8C,E and Supplementary Tables 6-7), with sufficient local resolution to confirm the absence of nucleotide in γ-tubulin*_Ce_* in an independent specimen (Supplementary Figure 8C,F). Supporting this result, addition of excess GTP (2 mM) prior to grid freezing successfully restored density for nucleotide in γ-tubulin*_Ce_*:GCP2*_Ce_*, as judged by a separate 3D cryo-EM reconstruction generated at 2.9 Å resolution from ∼400,000 particles (FSC_0.143_; Figure 4C, Supplementary Figure 8D-E and Supplementary Tables 6-7). This suggests that γ-tubulin*_Ce_* binds GTP with a lower affinity than, e.g., human γ-tubulin, which has a reported K_D_ of ∼50 nM (*27*, *48*). Notably, the single-spoked GTP-bound γ-tubulin*_Ce_*model also recapitulates the nucleotide-induced remodeling of helices H1, H2, H3’, and H5 when compared with other nucleotide-bound α-, β- or γ-tubulin structures (Figure 4D-E, Supplementary Figure 7D-E and Supplementary Video 1), suggesting that stabilization of these secondary structure elements likely correlates with nucleotide binding in all eukaryotic tubulins.

### 4-spoked assemblies oligomerize into microtubule-nucleating templates with 13-fold symmetry

Surprisingly, when processing the 4-spoked assembly cryo-EM data (Figure 3C), we also identified particles corresponding to 8-spoked (∼25% of particles), 12-spoked (∼14% of particles), and likely larger assemblies (Figure 5A and Supplementary Figure 6D; see also Methods). After supervised 3D classification of all starting 4-spoked assembly particles, we were able to generate 3D reconstructions at 3.3 Å resolution for the 8-spoked assembly and 5.9 Å resolution for the 12-spoked assembly (FSC_0.143_; Figure 5B, Supplementary Figures 6D, S9A-B, and Supplementary Tables 4-5). These resolutions allowed unambiguous assignment of the pseudo-helical complexes as head-to-tail multimers of the GCP2*_Ce_*/GTAP-1/GCP2*_Ce_*/GTAP-2-containing 4-spoked assembly found in γ-TuRC*_Ce_* (Figures 1C and 3C), with a now well-resolved ∼15 residue extension of the N-terminal portion of GTAP-1 stabilizing the multimerization interface (Figure 3D and Supplementary Figure 9C).

**Figure 5.**
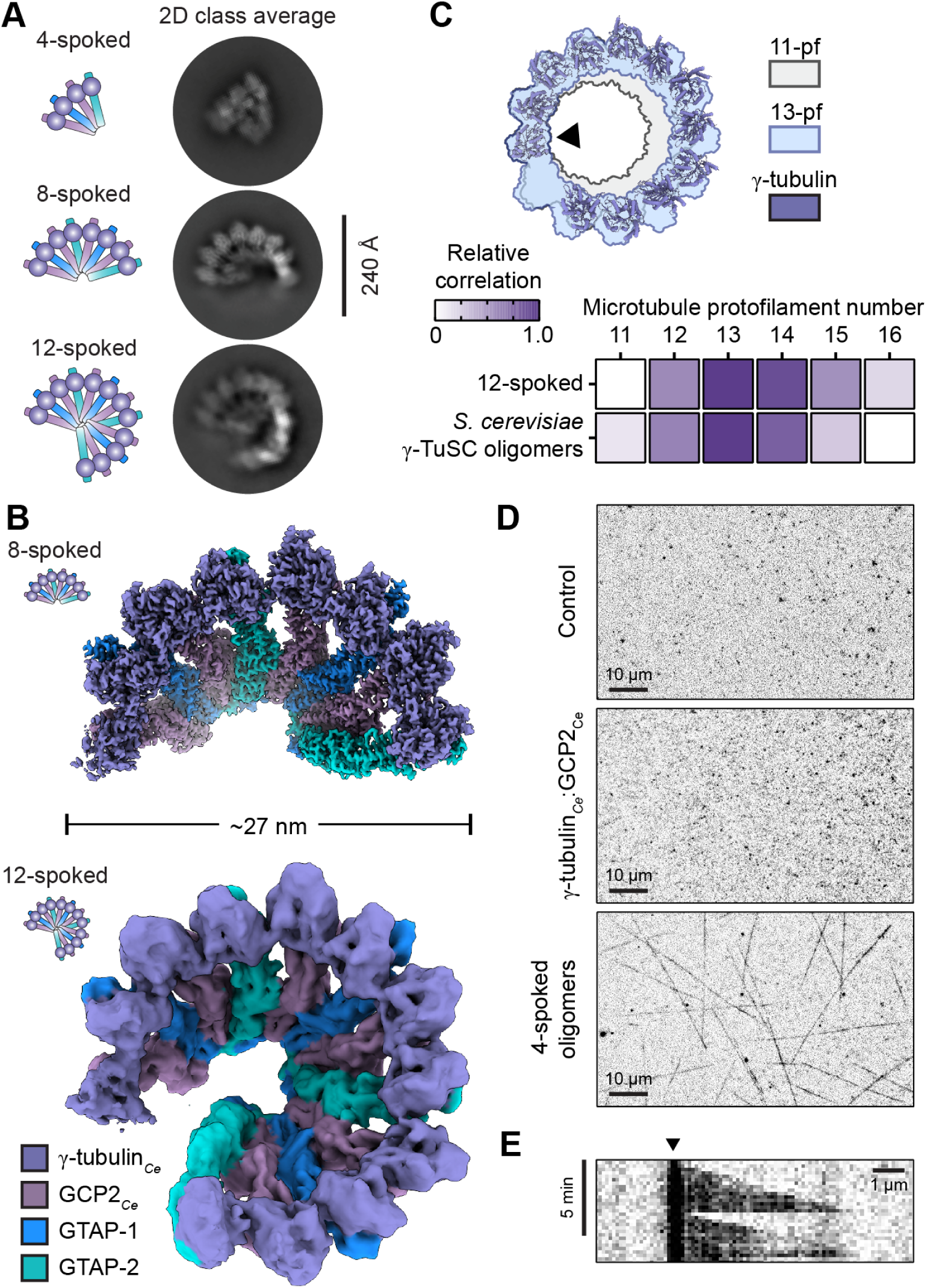
4-spoked assemblies oligomerize into microtubule-nucleating templates with ∼13-fold symmetry. **A)** Representative 2D class averages showing 4-spoked (top), 8-spoked (middle), and 12-spoked (bottom) assemblies identified by cryo-EM. Schematics of each assembly are shown on the left. **B)** Top views of cryo-EM densities for the 8-spoked assembly (top) and the 12-spoked assembly (bottom). Densities are colored according to the legend at the bottom left. The 8-spoked assembly is a composite map from multi-body refinements postprocessed using EMReady prior to combining (*44*). **C)** Top: γ-tubulin*_Ce_* ring from the 12-spoked assembly model aligned to a ring of β-tubulin from an 11-protofilament microtubule (grey; EMD-5191) or a 13-protofilament microtubule (light blue; EMD-5193) (*38*). The γ- and β-tubulin subunits used for alignment are indicated with a black triangle. Bottom: Cross-correlation values of β-tubulin rings from various microtubule protofilament number architectures fitted to the γ-tubulin rings from the 12-spoked assembly (this study) or budding yeast γ-TuSC oligomers in the “closed” conformation (PDB ID: 5FLZ (*39*)) as controls, colored according to the legend on the left (see Methods). **D)** TIRF microscopy fields of view of microtubule nucleation reactions containing tubulin alone (top; control), tubulin plus γ-tubulin*_Ce_*:GCP2*_Ce_*subcomplexes (middle), or tubulin plus reconstituted 4-spoked oligomeric assemblies (bottom). The tubulin concentration in all experiments was 10 μM. Three independent experiments were performed for each condition with similar results. **E)** Kymograph of a microtubule nucleated by 4-spoked oligomeric assemblies at a tubulin concentration of 10 μM tubulin. Black triangle indicates the inferred position of the nucleating site. The mean microtubule growth rate, measured from n = 10 microtubules from N = 3 experiments, was 3.8 +/- 0.9 μm/min.

In hindsight, multimerization of the 4-spoked assembly might have been anticipated, as the multimer interface comprises GTAP-2 from one 4-spoked assembly and GCP2*_Ce_* from the next (Figure 5A-B); after all, the same interface forms between GTAP-2 (position 10) and GCP2*_Ce_*(position 11) in γ-TuRC*_Ce_* (Figure 1C-D). In contrast with γ-TuRC*_Ce_*, however, the pseudo-helical multimeric 8- and 12-spoked assembly structures are no longer consistent with an ∼11-protofilament microtubule architecture. Instead, cross-correlation and rotational averaging analyses indicated that they now best match the canonical 13-protofilament α/β-tubulin lattice found in cytoplasmic microtubules of other species (Figure 5C and Supplementary Figure 9D-E). Remarkably, the reconstituted 4-spoked assemblies nucleated and capped the minus ends of porcine brain microtubules in TIRF microscopy assays (Figure 5D-E), suggesting they can function - either alone or as multimers - as γ-TuRC-like microtubule nucleation templates. Together, our results indicate that γ-tubulin*_Ce_* complexes exhibit an evolutionary adaptation that allows them to form microtubule-nucleating assemblies that can switch between ∼11- or ∼13-protofilament-like geometries.

## Discussion

In this study, we determined the structure of the native *C. elegans* γ-TuRC. This “γ-TuRC*_Ce_*” has previously been proposed to be built solely from repeating γ-TuSCs (*30*, *32*), similar to complexes in the fungi *S. cerevisiae* and *C. albicans* (*42*, *43*). We have shown that in addition to multiple γ-TuSCs, γ-TuRC*_Ce_* is additionally built from the nematode-specific GCP-like proteins GTAP-1 and GTAP-2. As γ-TuRC*_Ce_* was isolated from whole animals, GTAP-1 and GTAP-2 are likely core components of the complex, rather than part of germline-specific assemblies (*31*). Moreover, several lines of evidence suggest that these proteins most likely evolved from GCP5 (for GTAP-1) and GCP6 (for GTAP-2): i) their positioning in γ-TuRC*_Ce_* is analogous to the GCP5 and GCP6 subunits in vertebrate γ-TuRCs (*20*, *22*, *23*); ii) our bioinformatics analysis identified them as likely GCP5 and GCP6 orthologs (Supplementary Figure 5A-E); and iii) GTAP-2-NTD interacts with actin*_Ce_* via similar structural elements as GCP6-NTD in the vertebrate γ-TuRC (Figure 2B-C and Supplementary Figure 4B-C) (*21*). Unlike vertebrate GCP5 and GCP6, however, which are paired with GCP4 (*20*, *22*), GTAP-1 and GTAP-2 are paired with GCP2*_Ce_*, which is also a component of the γ-TuSCs occupying γ-TuRC*_Ce_* positions 3-6 and 11-12 (and, presumably, positions 1-2; Figure 1B-D and Supplementary Figure 3C). As GCP4 evolutionarily predates the emergence of nematodes (*49*) (Supplementary Figure 5A-D), GCP2*_Ce_* has likely co-evolved with GTAP-1 and GTAP-2 to accommodate the absent GCP4-like interfaces within positions 7-10 of γ-TuRC*_Ce_*, revealing an unexpected degree of co-evolution and interchangeability in the GCP family of proteins.

We propose that the lateral interfaces between the GRIP1 domains of *C. elegans* GCP subunits contribute to γ-TuRC*_Ce_*’s unusual, ∼11-fold symmetry. This is supported by: i) diameter variations between structures of γ-TuRC*_Ce_* and γ-TuRCs from other species, despite being built from GCPs with relatively conserved folds (Figure 1C-D) (*31*); ii) recapitulation of relative architectural differences between γ-TuRC*_Ce_* and other γ-tubulin subcomplexes by AlphaFold predictions (Supplementary Figure 10A-D); and iii) the fact that, during γ-tubulin ring closure, lateral GRIP1-GRIP1 interfaces in other γ-TuRCs remain relatively immobile even as GRIP2 domains undergo dramatic structural rearrangements (*36*, *39*, *50*). Consequently, our work suggests two distinct “control settings” within the GCPs that can tune overall γ-TuRC geometry. One is the arrangement of lateral GRIP1-GRIP1 interfaces, whose large buried surface areas dictate the number of GCPs allowed per helical turn (e.g., ∼11 vs. ∼13). The other is the GRIP1-GRIP2 inter-domain angles, which control the local orientation of γ-tubulin and the subsequent degree of constriction of the γ-tubulin ring (*36*). Our work shows that GCPs do more than facilitate microtubule nucleation, minus end capping, and γ-TuRC accessory protein recruitment; they also function as fundamental determinants of microtubule geometry.

The unique organization of actin*_Ce_* within γ-TuRC*_Ce_* may have consequences for the assembly and stability of the *C. elegans* complex. In vertebrate γ-TuRCs, actin makes two primary contacts with: i) GCP6-NTD, via the barbed end groove (*21*); and ii) the γ-tubulin:GCP3 interface at position 2, via the actin “D-loop” (*20*, *22*). In our γ-TuRC*_Ce_* cryo-EM reconstruction, the γ-tubulin*_Ce_*:actin*_Ce_*D-loop interface is absent because subunits at positions 1 and 2 are missing (Figure 1C-D). However, actin*_Ce_*’s polymerization interface, which is solvent-exposed in the vertebrate γ-TuRC, forms a buried interface with the base of the GRIP1 domains of GCP2*_Ce_* and GCP3*_Ce_*at positions 11-12 (Figure 2A-B). Additionally, GTAP-2-NTD contacts not only actin*_Ce_* but also the GRIP1 domain of GCP2*_Ce_* at position 3 (Figures 1C-D and 2B). Notably, γ-TuRC*_Ce_* lacks the stabilizing features seen in vertebrate γ-TuRCs, such as the “GCP6 belt” that spans multiple γ-TuSCs, or the MZT1:GCP5-NTD latch at the γ-TuRC seam (*51*). The presence of actin*_Ce_* in the cryo-EM reconstruction despite a missing D-loop interface, together with the absence of γ-TuSCs before position 1 and beyond position 12, suggests that actin*_Ce_* may regulate the incorporation of γ-TuSCs into γ-TuRC*_Ce_*. Since we do not observe any structural clash between actin*_Ce_* and additional γ-TuSCs in γ-TuRC*_Ce_* (not shown), actin*_Ce_*is unlikely to be an assembly “blocking” factor. Instead, we propose that an intrinsically higher rate of γ-TuSC disassembly from γ-TuRC*_Ce_* might be reduced through stabilizing contacts with: i) the terminal γ-TuSC via actin*_Ce_*’s polymerization interface; ii) the γ-TuSC at positions 3-4 via GTAP-2-NTD; and/or iii) the γ-TuSC at positions 1-2 via actin*_Ce_*’s D-loop. Differences in binding affinities across these interfaces could explain why the γ-TuSC at positions 1-2 is less stably incorporated than the one at positions 11-12. While the role of actin in γ-TuRCs remains an open question (*52*), our study indicates that divergent γ-tubulin complexes may offer a useful tool for dissecting its function in the future.

The NTD of GTAP-2 performs a similar actin-binding function as MZT1:GCP6-NTD in vertebrates (Figure 2). At our γ-TuRC*_Ce_* resolutions, we do not observe any densities for MZT1*_Ce_*bound to this domain nor to any other part of the complex (e.g. GCP3*_Ce_*), and we did not detect any MZT1*_Ce_* from MS analysis of γ-TuRC*_Ce_*(Supplementary Table 1). Our work suggests that in certain species the function of previously defined “MZT modules” such as actin-binding can be performed in a MZT protein-independent manner. This result may also help explain how γ-TuRCs can still assemble, and in some cases even function, in the absence of MZT proteins in a variety of species, including *C. elegans* (*30*), *D. melanogaster* (*53*), *S. pombe* (*54*), and in *C. albicans* (*43*).

The observation that γ-tubulin*_Ce_* is in the nucleotide-free (apo) state was unexpected for several reasons. First, our purifications were conducted in purification buffers containing standard GTP concentrations (10-100 μM) (*1*, *20*, *24*, *35*, *42*). Second, guanine nucleotides have been identified in all high-resolution γ-tubulin structures reported to date (see e.g., (*20*, *24*, *26*, *27*, *55*)). Lastly, the residues comprising the nucleotide binding pocket of γ-tubulin*_Ce_*are relatively well-conserved (Supplementary Figure 8H). To our knowledge, the apo state has only been captured in distantly related tubulin homologs such as FtsZ, TubZ, and the archaeal OdinTubulin (*56–58*). Our structure of apo-γ-tubulin*_Ce_* thus reveals a possible intermediate during the nucleotide exchange pathway of eukaryotic tubulins. Compared with GTP-bound γ-tubulin*_Ce_*, helices H1, H2, H3’, and H5 positioned in the vicinity of the nucleotide binding domain are disordered in apo-γ-tubulin*_Ce_*, while the rest of the tubulin core is largely intact (Figure 4D). We found similar differences between apo-γ-tubulin*_Ce_* and nucleotide-bound α- or β-tubulin (Figure 4B and Supplementary Figure 7A-E). This argues that nucleotide-dependent stabilization of the tubulin fold may be a common feature of all eukaryotic tubulins, and could also explain why tubulin can tolerate the relatively fast off-rate of GDP from β-tubulin (∼0.1 s^-1^) measured in the absence of nucleotide (*59*), potentially allowing for rapid nucleotide exchange in unpolymerized tubulin to take place in the absence of dedicated guanine nucleotide exchange factors.

γ-TuRC*_Ce_* adopts an ∼11-fold rotational symmetry that matches the 11-protofilament architecture of *C. elegans* cytoplasmic microtubules (*16*). This suggests that γ-TuRC*_Ce_*likely templates the non-canonical filaments in this organism. However, *C. elegans* also contain microtubule populations with non-11-protofilament architectures, including 13-protofilament centriolar A-microtubules (*15*, *16*). The presence of γ-TuRC-shaped caps at the ends of centriolar A-microtubules in human lymphoblast pro-centrioles (*60*), mouse ependymal cell basal bodies (*61*), and *Tetrahymena thermophila* basal bodies (*62*), all support a γ-tubulin-dependent mechanism for specifying the 13-protofilament architecture of the A-microtubule. Constriction of the γ-tubulin ring alone is likely insufficient to modify γ-TuRC*_Ce_*’s protofilament number from ∼11 to ∼13 and would anyway result in improperly-oriented γ-tubulin subunits (*36*). Therefore, a γ-TuRC-like assembly different from the one we have isolated from whole animals might instead be used to template centriolar A-microtubules in *C. elegans*. One solution could involve the GCP2/GTAP-1/GCP2/GTAP-2 4-spoked assembly, which we have identified as a stable subcomplex that can multimerize *in vitro* into a microtubule-nucleating template, and whose architecture is consistent with a 13-protofilament microtubule (Figure 5C and Supplementary Figure 9D-E). Given that GCP3*_Ce_* is essential in *C. elegans* (*30*), it is conceivable that incorporation of a single γ-TuSC to the 12-spoked assembly we have identified *in vitro* would generate a 14-spoked template with ∼13-fold radial symmetry (Supplementary Figure 9E), although further work will be needed to demonstrate the existence of such hypothetical complexes in cells. Nevertheless, our molecular characterization of microtubule-nucleating complexes in *C. elegans* suggests that γ-TuRC architecture and composition can be modified across evolutionary lineages - and potentially within the same cell type - to accommodate multiple microtubule lattice architectures.

## Acknowledgements

We thank Miroslav Peterek and Bilal Qureshi from ScopeM (ETH Zürich), as well as Tamino Cairoli and Pavel Afanasyev from the Cryo-EM Knowledge Hub (ETH Zürich) for cryo-EM sample preparation, data collection and data processing support. We thank Luca Valánszki for insect cell culture support. We thank Christian Landolt, Alex Myczko, and Emil Zylis (ETH Zürich) for IT support, as well as Amol Aher, Paul Conduit, Martin Pilhofer, and Sami Chaaban for feedback on early manuscript drafts. We thank Kuo-Chiang Hsia for sharing the MZT1:GCP6-NTD expression plasmids. This study includes calculations performed on the Euler cluster of ETH Zürich. Proteomics experiments were performed at the Functional Genomics Center Zurich (FGCZ) of University of Zürich and ETH Zürich. This work was supported by an SNSF Project Grant (#310030_208120) and an SNSF Starting Grant (#TMSGI3_211309), both to MW, and by NIH R35GM133573 to SY.

## Author contributions

RK and MW designed the study. RK and BC purified proteins used. YX generated *C. elegans* strains used in this study. RK collected all EM data and processed negative stain EM data. RK, DZ, HMH, and MW performed cryo-EM processing. RK and MW built protein models. BC performed GST pull-down experiments. RK and BC performed bioinformatic analyses. RK and FM performed TIRF microscopy assays. RK and MW wrote the paper with contributions from all other authors. SY and MW supervised the study and obtained funding.

## Data Availability

Cryo-EM maps have been deposited in the EMDB and are publicly available as of the date of publication. Molecular models have been deposited in the PDB and are publicly available as of the date of publication. Accession numbers are listed in the Supplementary Tables. This paper analyzes existing, publicly available data. These accession numbers for the datasets are listed throughout the manuscript, where appropriate. Data reported in this paper will be shared by the lead contact upon reasonable request. Any additional information required to reanalyze the data reported in this paper is available from the lead contact upon reasonable request.

## Methods

### Gene editing by CRISPR/Cas9

CRISPR-based tagging of *C. elegans* γ-tubulin with 1xFLAG separated by an SSSS linker in (shy266[tbg-1::1xFLAG]) was performed according to single-stranded oligonucleotide donor protocols (*63*, *64*). In brief, 0.5 μL Cas9 (*S. pyogenes* Cas9 Nuclease V3, 10 μg/μL, IDT #1081058), 2 μg gRNA, 2 pmol ssDNA repair template with 30 bp homology arms (synthesized by IDT), 800 ng of pRF4[rol-6(su1006)] plasmid as a selection marker, and nuclease-free water were added to reach a total volume of 20 μL. The 20 bp NGG PAM sites were selected using Benchling (benchling.com) with a >55 on-target score and >50% GC content. The gRNAs were synthesized from DNA oligos (Millipore Sigma) using the EnGen sgRNA synthesis kit (NEB #E3322S) and subsequently purified by Monarch RNA cleanup kit (NEB #T2040). The injection mix was centrifuged at 13,000 rpm for 20 min and the supernatant was kept on ice and used for injection. F1 rollers were single-picked and genotyped to screen the correct editing events. The strain was outcrossed to N2 wild-type strain once before experiments. PAM site: AAAAGGACTATCTGACAAGA; repair template sequence: atcccgcgagaataaataataataaatCTActtgtcatcgtcatccttgtaatcTGACGATGAGGAAAGGCCCCTAGTGAGGTAATCTTTTTGTACGACCGCCTTGTACTCGTCCAGC.

### Purification of γ-TuRC_Ce_

Whole *C. elegans* shy266 animals were cultured in 4 L of S-medium (for 1 L: 5.85 g NaCl, 1 g K_2_HPO_4_, 6 g KH_2_PO_4_, 1 ml cholesterol (5 mg/ml in ethanol), H_2_O to 1 L, supplemented with 10 ml 1 M potassium citrate pH 6, 10 ml trace metals solution, 3 ml 1 M CaCl_2_, 3 ml 1 M MgSO_4_ (*65*)) supplemented with 8 g/L of *E. coli* strain HB101 as a food source. Cultures were grown at 18 °C with shaking at 180 rpm for 1-2 weeks. Cultures were harvested by centrifugation at 1,000 x *g*, snap-frozen in liquid nitrogen, and stored at −80°C until purification.

Harvested *C. elegans* pellets were resuspended in ice-cold lysis buffer containing 40 mM HEPES–KOH pH 7.5, 100 mM KCl, 5% (v/v) glycerol, 1% (v/v) Triton X-100, 2 mM MgCl_2_, 2 mM DTT, 10 µM GTP, 10 µM ATP, 0.05% (v/v) Tween-20, Complete™ protease inhibitor cocktail (Roche), and Benzonase® (Merck; added according to the manufacturer’s instructions). Animals were lysed using an EmulsiFlex-C5 high-pressure homogenizer (Avestin) and clarified by ultracentrifugation at 185,000 x *g* for 30 min. The clarified supernatant was incubated for 2 hr with anti-FLAG M2 magnetic agarose beads (Merck), followed by magnetic separation using a neodymium magnet. Beads were washed five times with at least 20 resin volumes of wash buffer (identical to lysis buffer but without Triton X-100), and proteins were eluted with 2 resin volumes of 250 µg/ml FLAG peptide (Merck) in 20 mM HEPES–KOH pH 7.5, 80 mM KCl, 2 mM MgCl_2_, 2 mM DTT, and 100 µM GTP applied for 2 h on ice with occasional gentle manual bead resuspension. Freshly eluted material was used to prepare cryo-EM grids, whereas frozen aliquots were used for LC–MS/MS peptide identification and TIRF microscopy-based microtubule nucleation assays.

### Expression and purification of recombinant 4-spoked assemblies

Coding sequences for γ-tubulin*_Ce_* and MZT1*_Ce_* were polymerase chain reaction (PCR)-amplified from cDNA (Horizon *C. elegans* ORF Collection) and inserted into the NotI and XbaI sites of pACEBac1 via restriction ligation cloning. The γ-tubulin sequence was mutated via Q5 site directed mutagenesis (NEB) to remove unwanted BstXI sites, generating pACEBac1:γ-tubulin*_Ce_* and pACEBac1:MZT1*_Ce_*.

A fragment containing the coding sequence for GCP2*_Ce_*was codon optimized for *S. frugiperda* expression, synthesized (Twist Biosciences) and cloned into the HindIII and EcoRI sites of pACEBac1 via restriction ligation cloning. A TEV-cleavable ZZ-tag sequence was excised from a previously-described MZT2A construct (*35*), and inserted in frame with the C-terminus of GCP2 in pACEBac1 using restriction ligation cloning via EcoRI, KpnI, and HindIII sites, generating pACEBac1:GCP2*_Ce_*-TEV-ZZ.

A fragment containing the coding sequences for *C. elegans* GCP3 (gip-1b splice variant), GTAP-1, GTAP-2 were codon optimized for expression in *S. frugiperda*, synthesized (GENEWIZ Germany GmbH) and cloned into separate pACEBac1 vectors using BamHI and EcoRI sites via restriction ligation cloning, generating pACEBac1:GCP3*_Ce_*, pACEBac1:GTAP-1, and pACEBac1:GTAP-2.

Fragments containing the coding sequence for residues 1-261 of SPD-5 (SPD-5_1-261_) and C. elegans PLK1 with a T194D mutation (Plk1_T194D_) (*32*, *66*, *67*), were codon optimized for *S. frugiperda* expression, synthesized (Twist Biosciences) and cloned into the EcoRI site of pACEBac1 via restriction ligation cloning, generating pACEBac1:SPD-5_1-261_ and pACEBac1:PLK1_T194D_.

Next, pACEBac1:γ-tubulin*_Ce_*, pACEBac1:MZT1*_Ce_*, pACEBac1:GCP2*_Ce_*-TEV-ZZ, pACEBac1:GCP3*_Ce_*, and pACEBac1:SPD-5_1-261_ were sequentially assembled into a single polycistronic MutliBac donor plasmid using BstXI and I-CeuI sites (*68*), generating pACEBac1:γ-tubulin*_Ce_*/MZT1*_Ce_*/GCP2*_Ce_*-TEV-ZZ/GCP3*_Ce_*/SPD-5_1-261_.

Both MultiBac and standalone PLK1_T194D_, GTAP-1 and GTAP-2 genes in pACEBac1 were transformed into DH10MultiBacTurbo *E. coli* cells (ATG:biosynthetics GmbH) and subjected to blue-white screening for transposition-positive colonies. Positive colonies were grown overnight in 6 ml LB supplemented with gentamicin, kanamycin and tetracycline at 37°C, 180 rpm to amplify recombinant bacmids. Bacmids were purified using resuspension, lysis and neutralisation buffers from the GeneJET Plasmid Miniprep Kit (ThermoFisher) followed by three rounds of extraction with ice-cold UltraPure™ phenol:chloroform:isoamyl alcohol (25:24:1 (v/v); Invitrogen) with a final round of chloroform clean-up. The resulting water phase was supplemented with 0.8 M LiCl and precipitated in 80% EtOH at 4°C for 30 minutes, washed in ice-cold pure EtOH and left to dry at 37°C for 15 minutes. The resulting bacmid-containing DNA pellets were resuspended in DNAse-free sterile H_2_O for concentration measurement. For transfection, 25 µg of bacmid was mixed with 100 µg of polyethyleneimine (PEI Max) and used to transfect 25 ml of 1×10^6^ cells/ml of Sf9 cells in Sf-900™ III serum-free madia. All baculoviruses were propagated for no longer than three passages. For expression, 2 L of Sf9 cells at 2×10^6^ cells/ml were infected with 200 ml of P3 virus for the polycistronic MultiBac and supplemented with 100 ml of P3 of GTAP-1- and GTAP-2-encoding baculoviruses. The addition of 100 ml of P3 PLK1_T194D_ baculovirus was dispensable for complex assembly.

Cells were harvested by centrifugation at 1,000 x *g*, resuspended in 100 ml ice-cold lysis buffer (40 mM HEPES-KOH pH 7.5, 150 mM KCl, 2 mM MgCl_2_, 2% (v/v) glycerol, 2 mM DTT, 1 tablet Pierce Protease Inhibitor per 100 mL buffer, 0.05% (v/v) Tween 20, 10 µM GTP and 10 µM ATP), and lysed by dounce homogenization on ice. The lysate was cleared by centrifugation at ∼180,000 x *g* for 30 minutes, followed by binding to 0.4 ml of IgG Sepharose 6 Fast Flow affinity resin (Cytiva) for 2 hours on a rotary shaker at 4°C. Resin was washed with 100 ml of lysis buffer, after which a sample was proteolytically eluted for 2 hours with 1 mg of home-made TEV protease (from pRK793, a gift from David Waugh, Addgene plasmid 8827) resuspended in lysis buffer. All purification steps were analyzed by SDS-PAGE. Next, the TEV-eluate was clarified by centrifugation for 10 min at 17,000 x *g* at 4°C and separated using size exclusion chromatography (SEC) on a Superdex 200 Increase 10/300 GL column (Cytiva) pre-equilibrated with SEC buffer (20 mM HEPES-KOH pH 7.5, 150 mM KCl, 2 mM MgCl_2_, 2 mM DTT, 10 µM GTP). Fractions containing co-eluting GTAP-1, GTAP-2, GCP2*_Ce_* and γ-tubulin*_Ce_* were identified using SDS-PAGE, pooled together and concentrated on 35 kDa cut-off spin membrane (Millipore). Concentrated protein was aliquoted, snap-frozen in liquid nitrogen and stored at −80°C.

### Expression and purification of recombinant γ-tubulin_Ce_:GCP2_Ce_ subcomplexes

Expression and purification of the recombinant γ-tubulin*_Ce_*:GCP2*_Ce_*subcomplex was carried out in an identical way to that of 4-spoked assemblies but without the addition of GTAP-1 and GTAP-2 baculoviruses.

### Purification of MZT1:GCP6-NTD

Plasmids coding for GST-tagged GCP6-NTD and 6xHis-tagged MZT1 (a kind gift from Dr. Kuo-Chiang Hsia; (*21*)) were co-transformed into BL21 pRARE cells. Proteins were induced with 0.5 mM IPTG and expressed for 16 hr at 180 rpm and at 18°C. Cells were harvested by centrifugation at 5,000 x *g* for 20 minutes and at 4°C. Cell pellets were resuspended in Ni-NTA lysis buffer (50 mM HEPES pH 7.5, 200 mM NaCl, 2 mM MgCl_2_, 5% (v/v) glycerol, 2 mM 2-mercaptoethanol, 10 mM imidazole, supplemented with 1 protease inhibitor tablet per 150 ml buffer (Roche)) and lysed by 3 passes of an EmulsiFlex C5 microfluidizer (Avestin). Lysate was centrifuged at 50,000 x *g* for 40 min at 4°C and 0.2 μm filtered. The clarified lysate was then passed over a 5 ml HisTrap column (Cytiva) pre-equilibrated in Ni-NTA lysis buffer. The column was washed with 5 column volumes of Ni-NTA wash buffer (50 mM HEPES pH 7.5, 200 mM NaCl, 2 mM MgCl_2_, 5% (v/v) glycerol, 2 mM 2-mercaptoethanol, 20 mM imidazole, supplemented with 1 protease inhibitor tablet per 150 ml buffer (Roche)). Bound proteins were eluted with Ni-NTA elution buffer (50 mM HEPES pH 7.5, 200 mM NaCl, 2 mM MgCl_2_, 5% (v/v) glycerol, 2 mM 2-mercaptoethanol, 250 mM imidazole, supplemented with 1 protease inhibitor tablet per 150 ml buffer (Roche)). Eluted material was passed over a 5 ml GSTrap column (Cytiva) pre-equilibrated with GST wash buffer (50 mM HEPES pH 7.5, 200 mM NaCl, 2 mM MgCl_2_, 5% (v/v) glycerol, 2 mM 2-mercaptoethanol, supplemented with 1 protease inhibitor tablet per 150 ml buffer (Roche)). The column was washed with 5 column volumes of GST wash buffer, and bound proteins were eluted with GST elution buffer (50 mM HEPES pH 7.5, 200 mM NaCl, 2 mM MgCl_2_, 5% (v/v) glycerol, 2 mM 2-mercaptoethanol, 20 mM reduced glutathione). Protein-containing fractions were identified by Bradford assay, pooled, dialyzed against storage buffer (50 mM HEPES pH 7.5, 200 mM NaCl, 2 mM MgCl_2_, 5% (v/v) glycerol, 2 mM 2-mercaptoethanol) to remove excess glutathione, snap-frozen in liquid nitrogen, and stored at −80°C until use. Purified protein was analyzed by SDS-PAGE followed by Coomassie staining (Supplementary Figure 4D).

### Purification of GTAP-2-NTD

A construct expressing GTAP-2-NTD (residues 2 to 161) fused to an N-terminal 6xHis-GST-tag separated by an SGSGGGGG-TEV linker sequence was synthesized and cloned into a pET M30 vector (Azenta). GTAP-2-NTD was expressed and purified identically to MZT1:GCP6-NTD above. Purified protein was analyzed by SDS-PAGE followed by Coomassie staining (Supplementary Figure 4D).

### GST pulldown assays

For each reaction, 20 μL of Glutathione Sepharose 4B resin was equilibrated in pull-down buffer (50 mM HEPES pH 7.5, 200 mM NaCl, 2 mM MgCl2, 5% (v/v) glycerol, 2 mM 2-mercaptoethanol). 10 μL of GST, GST-tagged GTAP-2-NTD, or GST-tagged MZT1:GCP6-NTD (stock concentration 23 μM) was mixed with resin, and the reaction was diluted to 500 μL with storage buffer and incubated for 1 hr at 4°C with gentle nutation. The resin was centrifuged at 500 x *g* for 1 minute at 4°C and the supernatant was discarded. 10 μL of actin at 23 μM was added to the resin, and the reaction was again diluted to 500 μL with storage buffer and incubated for 1 hr at 4°C with gentle nutation. The resin was centrifuged at 500 x *g* for 1 minute at 4°C and the supernatant was discarded. The resin was washed three times with 500 μL of storage buffer followed by centrifugation. After the final wash, 2X SDS loading buffer was added directly to the pelleted resin and the sample was boiled at 95°C for 5 min. The resin was centrifuged a final time and the supernatant was resolved by SDS-PAGE for Coomassie staining or for western blotting using an anti-actin antibody (AAN02, Cytoskeleton).

### LC-MS/MS sample preparation, data acquisition and analysis

Samples were subjected to Sp3-assisted protein capture and clean-up. Proteins were reduced and alkylated by adding Tris(2-carboxyethyl)phosphine and 2-Chloroacetamide to a final concentration of 5 mM and 15 mM, respectively. The samples were incubated for 30 min at 30°C, 700 rpm and light-protected. Next, samples were diluted with pure ethanol to reach a final concentration of 60% (v/v). Using the KingFisher Flex System (Thermo Fisher Scientific) a corresponding amount of carboxylated magnetic beads (hydrophobic and hydrophilic) has been added to the samples. After binding the proteins to the beads for 30 min at room temperature, beads were washed 3x with 80% (v/v) EtOH. For the enzymatic digestion the beads were added to trypsin in 50 mM TEAB. Samples were digested overnight at 37°C. The remaining peptides were extracted from beads with H_2_O. The two elutions were combined and dried down. The digested samples were dissolved in aqueous 3% Acetonitrile with 0.1% formic acid, and the peptide concentration was estimated with the Lunatic UV/Vis absorbance spectrometer (Unchained Lab). Peptides were separated on a M-class UPLC and analysed on a Orbitrap mass spectrometer (Thermo). The acquired MS data were processed using the Maxquant search engine (V 2.0.1.0, (*69*)). The spectra were searched against the provided customer sequences merged with C. elegans UP000001940 protein background database. The following modifications were included in the analysis: Acetyl (Protein N-term), Oxidation (M), Deamidation (NQ), Carbamidomethyl (C). Ultimately, protein identification results have been imported and analysed in the Scaffold software (Proteome Software) using protein FDR of 1%, minimum 2 peptides required for a protein, and peptide FDR of 0.1%.

### Negative stain EM of γ-TuRC_Ce_

For negative stain transmission electron microscopy (TEM) analysis, carbon-coated copper grids (EMS; CF400-Cu-50) were glow-discharged using a Ted Pella PELCO easiGlow system at 25 mA for 30 s. Subsequently, 5 μl of γ-TuRC_Ce_ sample was applied to a grid held with inverted tweezers and incubated for 1 min before excess sample was blotted using Whatman No. 1 filter paper. Grids were immediately washed with a 20 μl drop of distilled water. Negative staining was performed by three consecutive 30 s incubations in 20 μl drops of 2% (w/v) uranyl acetate. Excess stain was removed by manual blotting, and grids were air-dried for >24 h in a sealed container containing desiccant prior to imaging. All datasets were acquired on a Thermo Fisher Scientific Talos L120C microscope at the ScopeM facility at ETH Zurich operated at 120 kV and a nominal magnification of 57,000 X, corresponding to a calibrated pixel size of 2.5 Å/pixel. Data were collected with a defocus range of −3.0 to −1.5 μm. Serial acquisition mode was used to collect 11,206 micrographs across three independent sessions. Micrographs were imported into cryoSPARC for image processing. An initial set of 669 particles was manually picked, of which 462 particles were classified into γ-TuRC-like 2D classes. These classes were subsequently used for iterative template-based particle picking, yielding 14,918 particles that contributed to an *ab initio* reconstruction of γ-TuRC*_Ce_*. Following heterogeneous refinement-based cleanup and multiple rounds of homogeneous refinement, the resulting density map was used to generate templates for uniform, orientation-independent particle picking across all three datasets to increase particle numbers. Subsequent 2D classification of template-based picks yielded a final dataset of 94,684 particles. Homogeneous and local refinement of these particles resulted in a reconstruction at 20 Å resolution. Further 3D classification into four classes identified a 12-spoked class containing approximately 20% of particles, which refined to a final resolution of 19 Å following local refinement. The negative stain EM density map has been deposited to the EMDB (EMD-58823).

### Cryo-EM of γ-TuRC_Ce_

For cryo-EM analysis of the native complex, Quantifoil R2/2 300-mesh copper holey carbon grids were coated in-house with a 1-2 nm continuous carbon layer and glow-discharged using a PELCO easiGlow device at 15 mA for 15 s. The grids were then mounted in Vitrobot-compatible tweezers pre-equilibrated on ice. To increase particle density, 3.6 μL of γ-TuRC*_Ce_* was manually applied to the grid and incubated for 30 s before excess liquid was removed using Whatman filter paper. This step was repeated 5-8 times. Following the final sample application, the tweezers were transferred to a Vitrobot (Thermo Fisher Scientific). Grids were blotted for 2 s with a blot force of −17 at 100% relative humidity and 8 °C and were immediately plunge-frozen into a liquid ethane-propane mixture. Frozen grids were subsequently stored in liquid nitrogen until imaging.

A total of 54,926 movies were collected across 2 data-collection sessions on a Titan Krios G3i transmission electron microscope (Thermo Fisher Scientific) equipped with a field-emission gun, a BioQuantum energy filter (Gatan) operated with a 20 eV slit width, and a K3 direct electron detector (Gatan) operating in correlated double sampling (CDS) mode. Data were acquired at the ScopeM facility, ETH Zürich, Switzerland. Automated data collection was performed in EPU (Thermo Fisher Scientific) using the Faster Acquisition mode. Total electron exposure was maintained within the range of 40 e⁻/Å², with images collected at defocus values ranging from −0.8 to −2.6 μm. Movies were recorded as 40-frame TIFF stacks.

Movie frames were motion-corrected and dose-weighted, followed by contrast transfer function (CTF) estimation and exposure curation for good CTF fit parameters. All processing steps were performed in cryoSPARC (*70*), unless otherwise indicated (Supplementary Figure 2A). Particles were identified through multiple rounds of Topaz-based particle picking (*71*), using the trained model for recombinant 4-spoked assemblies for the first iteration (Supplementary Figure 6D), as other picking strategies failed to detect γ-TuRC*_Ce_* particles. Multiple rounds of particle cleanup via 2D classification and repicking with new Topaz models trained on the cleaned up particles, followed by a final duplicate particle removal (minimum spacing 200 Å) led to 79,434 particles for Dataset 1 and 40,717 particles for Dataset 2.

For Dataset 1, binned particles were extracted and subjected to homogeneous refinement with the γ-TuRC*_Ce_* negative stain EM reconstruction as a starting reference (Figure 1B); 3D classification using a mask around the entire complex to identify particles with good secondary structure features; then heterogeneous refinement using 4 γ-TuRC*_Ce_* references, generated in silico to contain spokes at positions 1-12, 3-12, 5-12, or 7-12, since we observed evidence for a variety of such subcomplexes during 2D classification (see Supplementary Figure 2B for an example). The best class corresponded to the positions 3-12 reference and was used for further processing.

For Dataset 2, binned particles were subjected to one additional round of 2D classification, from which 34,503 particles were subjected to homogeneous refinement using the γ-TuRC*_Ce_* negative stain EM reconstruction as a starting reference (Figure 1B).

Dataset 1 and 2 particles were re-extracted and re-centered according to their alignments at a binning of 2, combined and subjected to heterogeneous refinement using 4 γ-TuRC*_Ce_* references, containing spokes at positions 1-12, 3-12, 5-12, or 7-12 as above for Dataset 1. The best class corresponded to the positions 3-12 reference. These 21,493 particles were subjected to homogeneous refinement followed by local refinement yielding the final γ-TuRC*_Ce_* consensus map (Figure 1C).

For model building, AlphaFold 3 was first used to predict the following starting models (*41*): *C. elegans* γ-TuSC (2 copies of γ-tubulin*_Ce_*, 1 copy of full-length GCP2*_Ce_*, and 1 copy of GCP3*_Ce_*); and the actin bridge (1 copy of GTAP-2-NTD and 1 copy of actin*_Ce_*). The refined recombinant 4-spoked assembly model below was used for final modeling of positions 7-10 of γ-TuRC*_Ce_*.

Subcomplex models were rigid body-fitted into the consensus γ-TuRC*_Ce_* cryo-EM reconstruction using ChimeraX “fit in map” or in Coot (*72*), breaking up subunits into domains when necessary. For some positions with lower density quality, model building was guided by additional AlphaFold Multimer predictions containing subunits at the interfaces between the various subcomplexes. Subunit models were then manually inspected, rebuilt if necessary, and real-space refined in Coot. Lastly, the entire model was subjected to a final real-space refinement in PHENIX to refine intersubunit contacts (*73*).

### Cryo-EM of recombinant 4-spoked assemblies

For cryo-EM analysis of 4-spoked assemblies, Quantifoil R2/2 300-mesh copper holey carbon grids glow-discharged using a PELCO easiGlow device at 25 mA for 30 s. The grids were then mounted in Vitrobot-compatible tweezers pre-equilibrated on ice. 3.6 μL of purified, concentrated 4-spoked assembly sample wasapplied to the grid and incubated for 30 s before excess liquid was manually removed using Whatman filter paper. This adsorption step was repeated up to 5 times. Following the final sample application, the tweezers were transferred to a Vitrobot (Thermo Fisher Scientific). Grids were blotted for 2 s with a blot force of −17 at 100% relative humidity and 8 °C and were immediately plunge-frozen into a liquid ethane-propane mixture. Frozen grids were subsequently stored in liquid nitrogen until imaging.

A total of 22,730 movies were collected across 2 data-collection sessions on either a Titan Krios transmission electron microscope (Thermo Fisher Scientific) equipped with a field-emission gun, a BioQuantum energy filter (Gatan) operated with a 20 eV slit width, and a K2 direct electron detector (Gatan) operating in correlated double sampling (CDS) mode, or a Titan Krios G3i transmission electron microscope (Thermo Fisher Scientific) equipped with a field-emission gun, a BioQuantum energy filter (Gatan) operated with a 20 eV slit width, and a K3 direct electron detector (Gatan) operating in correlated double sampling (CDS) mode. Data were acquired at the ScopeM facility, ETH Zürich, Switzerland. Automated data collection was performed in EPU (Thermo Fisher Scientific) using the Faster Acquisition mode. Total electron exposure was maintained within the range of 31 e⁻/Å², with images collected at defocus values ranging from −0.8 to −2.6 μm. Movies were recorded as 40-frame TIFF stacks.

Movie frames were motion-corrected and dose-weighted, followed by contrast transfer function (CTF) estimation and exposure curation for good CTF fit parameters. All processing steps were performed in cryoSPARC (*70*), unless otherwise indicated (Supplementary Figure 6D). Dataset 1 was used solely to generate templates for automated picking, as well as ab initio 3D references. Particles corresponding to 4-spoked assemblies in a variety of multimeric states were picked using blob picking from Dataset 1 micrographs and cleaned up via multiple rounds of 2D classification to remove junk particles. Templates from this step were used for template or Topaz-based autopicking from Dataset 2 (described below). They were separately used for template picking from Dataset 1 followed by 2D classification for particle cleanup and ab initio reconstructions, generating several classes corresponding to initial references for 4- and 8-spoked assemblies to aid in heterogeneous classification-based sorting of multimeric species in Dataset 2 particles.

For Dataset 2, templates generated above were used for several rounds of either template-based automated picking with 2D classification-based particle cleanup resulting in 1,553,123 initial particles, or else for Topaz-based picking with 2D classification-based particle cleanup resulting in 902,778 initial particles. To select specifically for non-multimerized 4-spoked assembly particles, initial particles from both picking methods were subjected to separate heterogeneous refinements using 6 classes: a 4-, 8-, and 12-spoked assembly class, generated either from Dataset 1 ab initio reconstructions as mentioned, or from ab initio reconstructions during targeted 2D classification selecting for classes representing specific multimeric species, as well as 3 additional “decoy” classes generated via ab initio reconstruction of junk particles identified during initial 2D classifications. Particles corresponding to the 4-spoked assembly class were identified from both heterogeneous refinements and combined. Duplicate particles were removed (minimum spacing 100 Å) and re-extracted with a binning of 2 and with recentering, resulting in 558,453 4-spoked assembly particles. Particles were subjected to non-uniform refinement using one of the heterogeneous refinement results as a reference, followed by several rounds of local and CTF refinement, resulting in a consensus 4-spoked assembly reconstruction at 3.0 Å resolution. To improve the local quality of the map, particles were exported into RELION 5, subjected to 3D autorefine with restricted angular and translation shifts as well as Blush regularization. Refined particles were then subjected to Multibody refinement using two bodies covering either the first or last two spokes of the 4-spoked assembly. This resulted in reconstructions with improved local density quality at resolutions of 3.0 - 3.1 Å (Supplementary Figure 6D).

To generate the 8-spoked assembly reconstruction, particles from the heterogeneous refinement job above that used template-based picking as input particles corresponding to the 8-spoked assembly class were selected for further processing. Duplicate particles were removed (minimum spacing 100 Å), subjected to non-uniform refinement using the 8-spoked assembly heterogeneous refinement result as an initial reference, and re-extracted with a binning of 2 and with recentering, resulting in 331,663 8-spoked assembly particles. Particles were subjected to non-uniform, local, and CTF refinements before performing per-particle reference based motion correction. A final local refinement resulted in a consensus 8-spoked assembly reconstruction at 3.3 Å resolution. To improve the local quality of the map, particles were exported into RELION 5, subjected to 3D autorefine with restricted angular and translation shifts as well as Blush regularization. Refined particles were then subjected to Multibody refinement using four bodies covering each distinct two spoke “pair” within the 8-spoked assembly. This resulted in reconstructions with improved local density quality at resolutions of 3.2 - 3.4 Å (Supplementary Figure 6D).

Lastly, to generate the 12-spoked assembly reconstruction, particles from the heterogeneous refinement job above that used template-based picking as input particles corresponding to the 12-spoked assembly class were selected for further processing. Particles were subjected to non-uniform refinement using the 8-spoked assembly heterogeneous refinement result as an initial reference, local refinement, then 3D classification to select for the best-resolved 12-spoked assemblies, resulting in 20,361 particles. Particles were subjected to non-uniform refinement and re-extracted with a binning of 4 with recentering. A final local refinement resulted in a consensus 12-spoked assembly reconstruction at 5.9 Å resolution (Supplementary Figures 6D and 9A-B).

For model building of the 4-spoked assembly, AlphaFold 3 was first used to predict a starting model containing 4 copies of γ-tubulin*_Ce_*, 2 copies of full-length GCP2*_Ce_*, and 1 copy each of full-length GTAP-1 and GTAP-2 (*41*). A composite map was generated by fitting both multi-body refinement maps into the consensus map using ChimeraX “fit in map”, then combining using ChimeraX “vop max” (*74*). Subunits in the predicted 4-spoked assembly model were individually rigid body-fitted into the composite map using ChimeraX “fit in map” or in Coot (*72*), breaking up subunits into domains when necessary. Subunit models were manually inspected, rebuilt if necessary, and real-space refined in Coot. Lastly, the entire 4-spoked assembly model was subjected to a final real-space refinement in PHENIX to refine intersubunit contacts (*73*).

To build the 8-spoked assembly model, a composite map was first generated by fitting the four multi-body refinement maps into the 8-spoked assembly consensus map using ChimeraX “fit in map”, then combining using ChimeraX “vop max”. Two copies of the refined 4-spoked assembly model were rigid body-fitted into the composite map using ChimeraX “fit in map”. Each subunit was manually inspected, rebuilt if necessary, and real-space refined in Coot. An AlphaFold Multimer prediction containing only the GRIP1 domains of GCP2*_Ce_* (five copies), GCP3*_Ce_* (two copies), GTAP-1 and GTAP-2 helped guide model building of the extended GTAP-1 NTD across the interface between neighbouring 4-spoked assemblies. Lastly, the entire 8-spoked assembly model was subjected to a final real-space refinement in PHENIX to refine intersubunit contacts.

The 12-spoked assembly model was built by rigid body fitting individual refined spokes (e.g., γ-tubulin*_Ce_*:GCP2*_Ce_*) from the 4-spoked assembly into the 12-spoked assembly density map.

### Cryo-EM of recombinant γ-tubulin_Ce_:GCP2_Ce_ subcomplexes (apo & GTP-bound)

For cryo-EM analysis of the recombinant γ-tubulin*_Ce_*:GCP2*_Ce_*subcomplexes, 3.6 μL of purified protein at 1.5 mg/mL was applied to glow-discharged Quantifoil R2/2 300-mesh copper holey carbon grids. Grids were glow-discharged in a PELCO easiGlow device at 25 mA for 30 s and plunge-frozen using a Vitrobot Mark IV (Thermo Fisher Scientific). Samples were blotted for 2 s with a blot force of −17 at 100% relative humidity and 8 °C before immediate vitrification in a liquid ethane–propane mixture. Frozen grids were stored in liquid nitrogen until imaging. For γ-tubulin*_Ce_*:GCP2*_Ce_* in the GTP-bound state, samples were supplemented with 2 mM GTP prior to grid freezing.

For γ-tubulin*_Ce_*:GCP2*_Ce_* subcomplexes in the apo state, a total of 9,330 movies were collected on a Titan Krios G3i transmission electron microscope (Thermo Fisher Scientific) equipped with a field-emission gun, a BioQuantum energy filter (Gatan) operated with a 20 eV slit width, and a K3 direct electron detector (Gatan) operating in correlated double sampling (CDS) mode. Data were acquired at the ScopeM facility, ETH Zürich, Switzerland. Automated data collection was performed in EPU (Thermo Fisher Scientific) using the Faster Acquisition mode. Total electron exposure was maintained within the range of 59 e⁻/Å², with images collected at defocus values ranging from −0.5 to −2.5 μm. Movies were recorded as 40-frame TIFF stacks.

Movie frames were motion-corrected and dose-weighted, followed by contrast transfer function (CTF) estimation and exposure curation for good CTF fit parameters. All processing steps were performed in cryoSPARC (*70*), unless otherwise indicated (Supplementary Figure 8C). Particles were initially identified via “blob” picking followed by 2D classification and cleanup. Classes corresponding to γ-tubulin*_Ce_*:GCP2*_Ce_*-shaped subcomplexes were selected and used as references for template picking. Subsequent 2D classification and cleanup resulted in 109,344 particles. These particles were used for ab initio 3D reference generation with 3 classes, resulting in one reasonable γ-tubulin*_Ce_*:GCP2*_Ce_* subcomplex-shaped density and two “junk” references. The references were used for further particle cleanup via heterogeneous refinement, resulting in 53,207 particles in the “good” class. These particles were then re-extracted at a binning of 2, subjected to non-uniform refinement followed by local refinement, resulting in a final 5.9 Å resolution reconstruction (Supplementary Figure 8C).

For γ-tubulin*_Ce_*:GCP2*_Ce_* subcomplexes in the GTP state, a total of 16,191 movies were collected on a Titan Krios G3i transmission electron microscope (Thermo Fisher Scientific) equipped with a field-emission gun, a BioQuantum energy filter (Gatan) operated with a 20 eV slit width, and a K3 direct electron detector (Gatan) operating in correlated double sampling (CDS) mode. Data were acquired at the ScopeM facility, ETH Zürich, Switzerland. Automated data collection was performed in EPU (Thermo Fisher Scientific) using the Faster Acquisition mode. Total electron exposure was maintained within the range of 30 e⁻/Å², with images collected at defocus values ranging from −0.6 to −2.6 μm. Movies were recorded as 40-frame TIFF stacks.

Movie frames were motion-corrected and dose-weighted, followed by contrast transfer function (CTF) estimation and exposure curation for good CTF fit parameters. All processing steps were performed in cryoSPARC (*70*), unless otherwise indicated (Supplementary Figure 8D). Particles were initially identified via template matching using the reconstruction of the γ-tubulin*_Ce_*:GCP2*_Ce_*subcomplex in the apo state above to generate reference templates, followed by 2D classification and cleanup. The resulting particles were subjected to three processing steps to maximize the number of particles recovered from the data.

1. Particles were subjected to heterogeneous refinement using the three ab initio references generate for γ-tubulin*_Ce_*:GCP2*_Ce_* subcomplexes in the apo state above, as well as two more “junk” classes corresponding to the junk reconstructions from the heterogeneous refinement of the apo dataset. This resulted in 241,535 particles in the “good” subcomplex class.
2. A 2D class corresponding to the γ-tubulin*_Ce_*:GCP2*_Ce_* subcomplex side view was selected and used for Topaz picking, followed by 2D classification and cleanup. Reasonable particles were then subjected to the same 5-class heterogeneous refinement as in strategy 1, resulting in 142,054 particles in the “good” subcomplex class.
3. A 2D class corresponding to the γ-tubulin*_Ce_*:GCP2*_Ce_* subcomplex top view was selected and used for Topaz picking, followed by 2D classification and cleanup. Reasonable particles were then subjected to the same 5-class heterogeneous refinement as in strategy 1, resulting in 211,571 particles in the “good” subcomplex class.

Particles from all three strategies were re-extracted and re-centered using their alignment shifts at a binning of 2 (1.3 Å/pixel) and combined. Duplicate particles were removed (minimum spacing 100 Å). Combined particles were then subjected to non-uniform, CTF, and local refinements, resulting in a reconstruction at 3.5 Å resolution. To improve the resolution, and because of local resolution deterioration especially towards the GRIP1 domain of GCP2*_Ce_* in the subcomplex, refined particles were exported into RELION 5, subjected to 3D autorefinement using Blush regularization (*75*), followed lastly by DynaMight reconstruction, resulting in a final reconstruction at 2.9 Å resolution (Supplementary Figure 8D).

For model building of the apo γ-tubulin*_Ce_*:GCP2*_Ce_*subcomplex, a single γ-tubulin*_Ce_*:GCP2*_Ce_* “spoke” from the 4-spoked assembly model above was used as a starting point. γ-tubulin*_Ce_*, the GRIP2 domain of GCP2*_Ce_*, and the GRIP1 domain of GCP2*_Ce_* were rigid body-fitted into the consensus apo γ-tubulin*_Ce_*:GCP2*_Ce_* subcomplex map using ChimeraX “fit in map”. The models were manually inspected, rebuilt if necessary, re-connected, and real-space refined in Coot. Lastly, the entire apo γ-tubulin*_Ce_*:GCP2*_Ce_* subcomplex model was subjected to a final real-space refinement in PHENIX to refine intersubunit contacts.

The same strategy was used for model building of the GTP-bound γ-tubulin*_Ce_*:GCP2*_Ce_*subcomplex, except that the starting model for γ-tubulin*_Ce_* was derived from an AlphaFold3 prediction containing a GTP molecule (*41*).

### Cross-correlation analysis

Models of laterally-associated, partial β-tubulin rings were generated by fitting multiple copies of PDB ID: 1TUB into 11-3, 12-3, 13-3, 14-3, 15-4, or 16-4 cryo-EM density maps (*38*, *46*) using ChimeraX’s “fit in map” command (*74*). α-tubulin subunits were deleted and maps consisting of 9 laterally-associated β-tubulin partial rings were generated with the “molmap” command in ChimeraX, setting the “resolution” to 5 Å. γ-tubulin rings from either γ-TuRC*_Ce_*(9 available subunits), recombinant 12-spoked assembly (12 available subunits), or other γ-TuRCs (14 available subunits) were converted into density maps using the same procedure as for β-tubulin. β-tubulin partial ring maps were sequentially fitted into γ-tubulin ring maps using the “fit in map” command, manually rotating the maps and repeating the fits such that all (up to 14) γ-tubulin ring positions were sampled by the partial β-tubulin ring. This was done to account for potential fitting errors and for local variations in the various γ-tubulin rings. The cross-correlation of the resulting fits was calculated using ChimeraX’s “measure correlation” command and averaged across all rotational fits, if applicable. The averages were then normalized between 0 and 1 for each γ-tubulin assembly and are reported in Figures 1F and 5C.

### Rotational averaging analysis

To estimate and compare the rotational symmetry between γ-TuRC assemblies, we first sought to generate projections along the semi-helical axis of each complex. For each γ-TuRC assembly, we aligned the structure of α-tubulin in the α/β-tubulin heterodimer (PDB ID: 1TUB; (*46*)) to each γ-tubulin subunit in the complex. A vector was determined between the centers of mass of each γ-tubulin and β-tubulin in its aligned α/β-tubulin heterodimer. A consensus vector for the γ-TuRC assembly was generated by averaging all the individual γ-tubulin:β-tubulin vectors. All γ-TuRC models were then aligned to one another via their consensus vectors and collectively rotated to make the consensus vectors all point along the z-axis. Models were then converted into density maps using the “molmap” command in ChimeraX, setting the “resolution” to 5 Å (*74*). Projections were generated for each aligned γ-TuRC using RELION-5.0’s “relion_project” command (*76*) and used for subsequent rotational averaging and power spectra analysis.

Analysis of the γ-TuRC assembly projections was performed using a custom Python pipeline. Projections were first centered using a mask-based centroid detection algorithm. Binary masks were generated via Otsu thresholding or adaptive Gaussian thresholding, followed by morphological closing and opening operations to remove noise. The image was then shifted to align the mask’s centroid with the center of the image frame using an affine transformation matrix. To quantify symmetry, the script calculates intensity profiles along the circumference of 10 concentric rings sampled linearly from the center toward the edge of the particle box. For each ring, intensity values are extracted at 1° radial increments using bilinear interpolation. A Fast Fourier Transform (FFT) is applied to each individual ring profile to compute its power spectrum. Individual power spectra are then averaged to generate a single “average power spectrum” for the entire assembly, representing the mean magnitude across all sampled radii. Values from the resulting average power spectrum ranging from 11- to 16-fold rotational symmetry are reported in Supplementary Figures 3E, 8E and 9D. For qualitative visualization, the script generates 11- to 16-fold rotational averages by iteratively rotating the centered images by 360° degrees divided by the protofilament number and computing the mean intensity across all orientations (Supplementary Figures 3D, 8D and 9C).

### Orthology analysis

For HMMER generation and gene assignment, available sequences from various species were collected for GCP2, GCP3, GCP4, GCP5, GCP6, GTAP-1 and GTAP-2, from the Oma Orthology database, selecting for a 1:1 “Relationship type” (https://omabrowser.org/) and/or OrthoDB. Sequence sets were aligned using MAFFT E-INS-i and HMMER profiles were built from the alignments using the “hmmbuild” command from HMMER version 3.4 (hmmer.org). All HMMER profiles were tested against a set of nematode proteomes and the top scoring hits recorded in a matrix file.

For species tree generation (Supplementary Figure 5A), 18S alignments for species were downloaded from the silva repository (https://www.arb-silva.de/). A phylogenetic tree representing species was built using RAxML with 1000 replicates. The species *Echinococcus canadensis, Spirometra erinaceieuropaei, Schistocephalus solidus, Dibothriocephalus latus* (*Platyhelminthes* phylum) was used as an outgroup. The resulting tree was visualised using iTOL (https://itol.embl.de).

For HMMER orthology detection (Supplementary Figure 5B,C,E), HMMER profiles for GTAP-1 and GTAP-2, GCP2, GCP3, GCP4, GCP5 and GCP6 were used to search against GCP/GTAP protein sets recorded in the gene assignment matrix. Top bitscore hits for each set were recorded.

### TIRF microscopy of γ-TuRC_Ce_

The TIRF microscope setup was built on a Nikon N-STORM microscope equipped with an SR Apochromat TIRF 100X immersion oil objective with a numerical aperture of 1.49, three-axis piezo-electric stages MCL NanoDrive PiezoZ Drive, a Hamamatsu Orca Flash 4 v3 sCMOS camera, and a laser generating 561 nm excitation light. The microscope was controlled using NIS Elements AR 5.21.01 (Nikon). Glass slides (Premiere 8201) and glass coverslips (Corning, 22 × 22 mm, 2845–22) were washed for 5 min in acetone, rinsed with deionized water, sonicated for 5-10 min in 50% methanol, rinsed with deionized water, sonicated for 5-10 minutes in 0.5 M KOH, rinsed with deionized water and dried using filtered compressed air. Clean, dry coverslips were subjected to plasma cleaning for 3 minutes at high intensity in a Harrick Plasma PDC-32G-2 plasma cleaner connected to an ICME M71B4 vacuum pump. Coverslips were made hydrophobic by silanization with 0.1% dichlorodimethylsilane (v/v) in heptane for 2 hr. After silanization, the coverslips were submerged in clean heptane and sonicated for 20 minutes, followed by a 10 minute sonication in ethanol. The coverslips were washed in deionized water and dried using filtered compressed air. Silanized coverslips were attached to clean glass slides using 3 strips of double-sided tape separated by ∼5 mm to create two channels per slide. On the day of each experiment, aliquots of Alexa Fluor 546-labeled (*77*) and porcine cycled tubulin (*78*) prepared in-house were thawed, mixed to a final tubulin labeling ratio of ∼10%, and centrifuged at 90,000 rpm for 10 min in a TLA 120.1 rotor (Beckman). The concentration as well as labeling ratio was measured using an IMPLEN NanoPhotometer N60. The tubulin was re-aliquoted and snap-frozen in liquid nitrogen until further use on the same day. For experiments in Supplementary Figure 1A-B, the channel was rinsed with BRB80 (80 mM K-PIPES, 1 mM K-EGTA, 1 mM MgCl2, pH 6.8), followed by monoclonal anti-FLAG M2 antibody (Sigma Aldrich) diluted 1:50 (v/v) in BRB80. After 5 min of incubation, non-adherent molecules were washed out with BRB80 and the channel was treated for 5 minutes with 1% Pluronic F127 (Sigma Aldrich) in BRB80. After washing with BRB80, γ-TuRC*_Ce_* or an additional wash with BRB80 for the no-γ-TuRC*_Ce_* control was introduced for 5 minutes. The channel was subsequently washed with an assay buffer containing BRB80 + 1 mM DTT, 50 mM KCl, 0.15% (w/v) methylcellulose (Sigma Aldrich), 0.2 mg/ml bovine serum albumin (BSA), and 1 mM GTP (Jena Bioscience). A reaction mix containing 12.5 μM Alexa Fluor 546-labeled tubulin at a ∼10% labeling ratio in assay buffer and supplemented with an oxygen scavenger system (freshly mixed 0.035 mg/ml catalase (Sigma Aldrich), 0.2 mg/ml glucose oxidase (Merck), 2.5 mM glucose (Sigma Aldrich), and 10 mM DTT) was introduced into the flow cell. The flow cell was immediately sealed with VALAP (1:1:1 vaseline, linoleum, paraffin wax) and placed on the stage of the TIRF microscope, which was equilibrated to 35 °C prior to measurement. Timelapses of 3-4 randomly selected fields of view were collected at 10-second intervals and with an excitation wavelength of 561 nm (<2.7 mW, 20 ms exposure). Time lapse images were acquired for up to 10 minutes. A total of 4 channels of γ-TuRC*_Ce_* were measured at different surface-immobilization concentrations, with 3-4 fields of view each; 2 control (no γ-TuRC*_Ce_*) channels were measured with 3-4 fields of view each. Image sequences were subsequently processed using drift correction to compensate for stage and sample movement, followed by background subtraction in Fiji (*79*). For visualization purposes, image contrast was inverted uniformly across datasets to facilitate the clear representation of microtubule nucleation events.

### TIRF microscopy of recombinant 4-spoked assemblies

Analysis of the microtubule nucleation capacity of the recombinant 4-spoked assemblies was performed using the same TIRF microscopy setup, glass functionalization procedure, and assay buffers as described for native γ-TuRC*_Ce_*. SEC-purified fractions containing 100 nM 4-spoked assemblies were introduced into the silanized flow chamber and allowed to adsorb to the hydrophobic surface. Unbound material was removed by washing with BRB80, followed by incubation with 1% (w/v) Pluronic^TM^ F-127 in BRB80 to minimize nonspecific surface interactions. After an additional wash with BRB80, a reaction mixture containing 10 μM mix of Alexa Fluor 546-labeled tubulin (Cytoskeleton) and porcine cycled tubulin (*78*) at a ∼10% labeling ratio in 1X assay buffer was introduced into the chamber. As a control, the recombinant single-spoked GCP2*_Ce_*:γ-tubulin*_Ce_*complex was immobilized and analyzed under identical conditions at a concentration of 400 nM, corresponding to an equimolar concentration of γ-tubulin molecules relative to the 4-spoked assemblies. Time-lapse image series were acquired over 10-15 minutes from 8-15 independent fields of view per condition. Image sequences were subsequently processed using drift correction to compensate for stage and sample movement, followed by background subtraction in Fiji (*79*). For visualization purposes, image contrast was inverted uniformly across datasets to facilitate the clear representation of microtubule nucleation events.

## Supplementary Figures

**Supplementary Figure 1.**
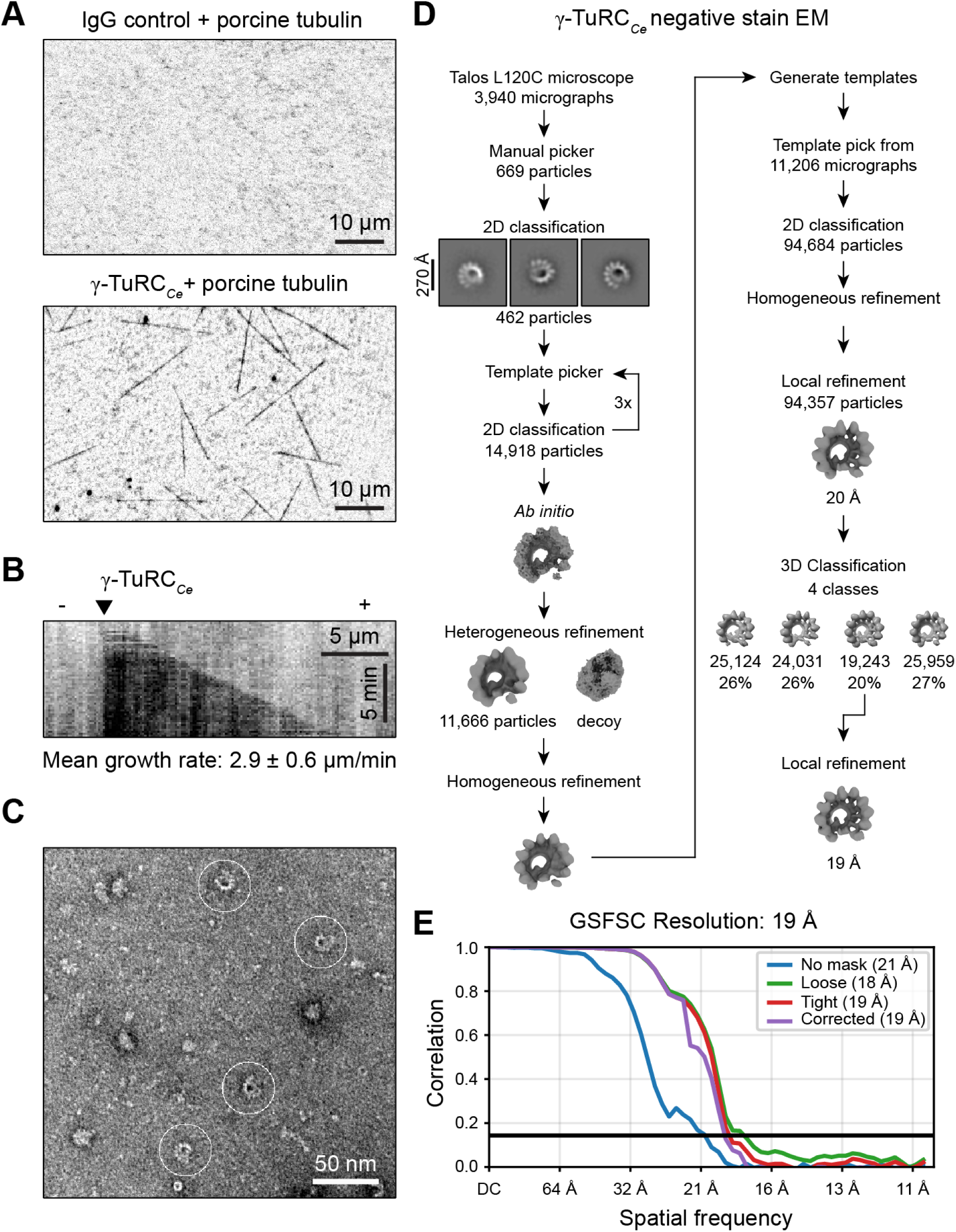
Characterization of γ-TuRC*_Ce_* by TIRF microscopy and negative stain EM. **A)** TIRF microscopy fields of view of microtubule nucleation reactions containing 12 μM fluorescently-labeled porcine brain tubulin without (top) or with (bottom) surface-immobilized γ-TuRC*_Ce_*. Three independent experiments were performed for each condition with similar results. **B)** Kymograph of a microtubule nucleated by γ-TuRC*_Ce_*, with the black triangle indicating the inferred position of the nucleating site. The mean microtubule growth rate measured from n = 12 microtubules from N = 3 experiments is indicated. **C)** Negative stain EM micrograph of isolated γ-TuRC*_Ce_*. Particles corresponding to γ-TuRC*_Ce_* are highlighted by white circles. **D)** SPA processing workflow for γ-TuRC*_Ce_* negative stain EM 3D reconstruction shown in Figure 1B. **E)** Gold standard FSC curve for the γ-TuRC*_Ce_* negative stain EM 3D reconstruction, generated in CryoSPARC (*70*).

**Supplementary Figure 2.**
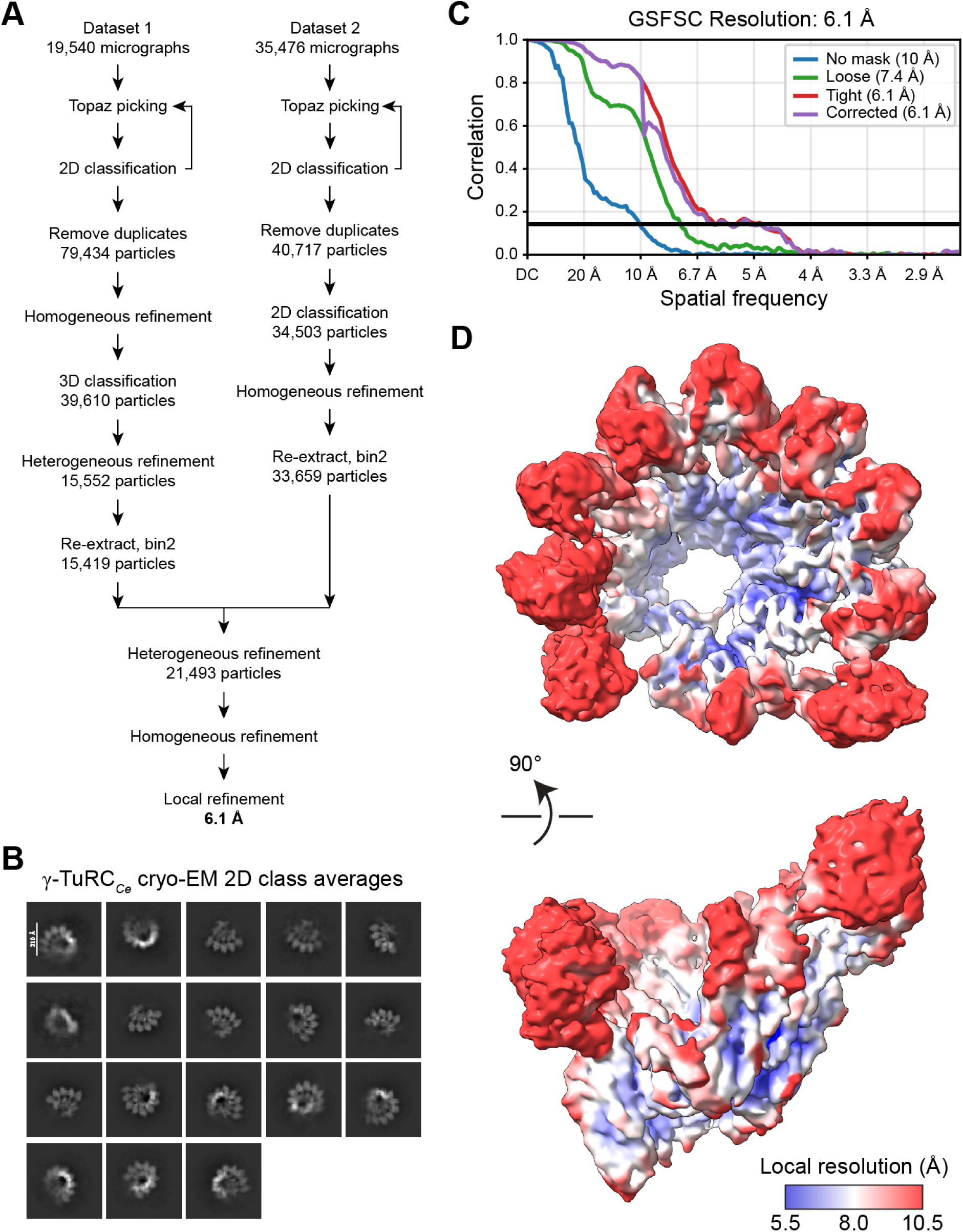
Processing details for the γ-TuRC*_Ce_* cryo-EM reconstruction. **A)** SPA processing workflow for γ-TuRC*_Ce_* cryo-EM 3D reconstruction shown in Figure 1C. **B)** 2D class averages of the combined, re-extracted bin2 particles at the end of each Dataset processing scheme in **A)**. **C)** Gold standard FSC curve for the γ-TuRC*_Ce_* cryo-EM 3D reconstruction, generated in CryoSPARC (*70*). **D)** Top and side views of γ-TuRC*_Ce_* cryo-EM reconstruction colored according to local resolution, calculated in CryoSPARC (*70*).

**Supplementary Figure 3.**
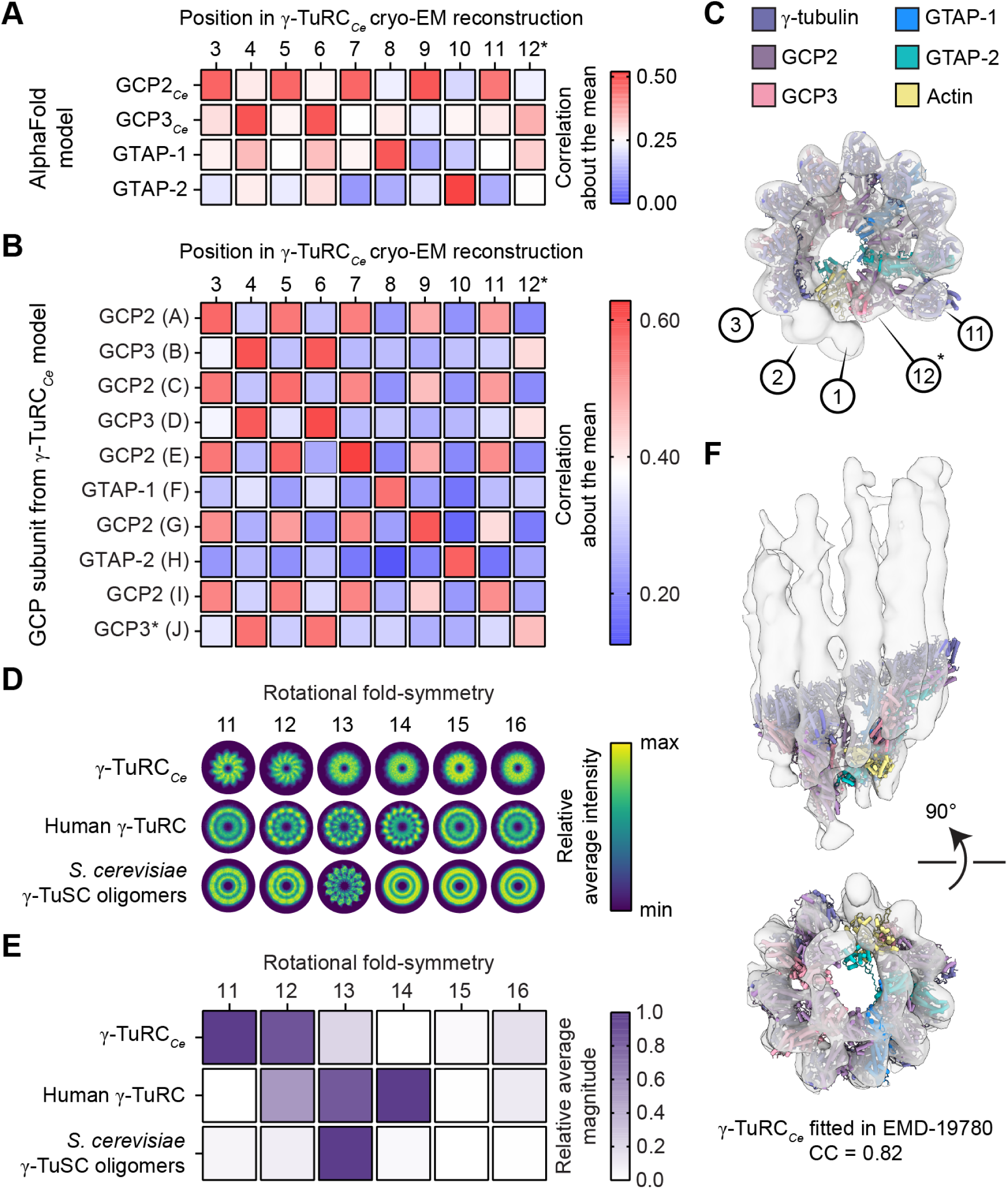
γ-TuRC*_Ce_* subunit identification and architecture analysis. **A)** Correlation-about-the-mean, as calculated in ChimeraX (*74*), for rigid body fits of GRIP1 domains of various GCP AlphaFold 3 models (y-axis) across the various positions of the γ-TuRC*_Ce_* cryo-EM reconstruction. **B)** Correlation-about-the-mean, as calculated in ChimeraX (*74*), for GRIP1 domains of each GCP subunit from the final γ-TuRC*_Ce_* model rigid body fitted into each possible position in the γ-TuRC*_Ce_* cryo-EM reconstruction. **C)** Top view of a cartoon representation of the cryo-EM derived γ-TuRC*_Ce_* model rigid body fitted into the γ-TuRC*_Ce_* negative stain EM reconstruction (transparent surface). γ-TuRC*_Ce_* subunits are colored according to the legend at the top. γ-TuRC*_Ce_* positions 1, 2, 3, 11, and 12 are indicated. Asterisk indicates the partially resolved spoke at γ-TuRC*_Ce_*position 12. **D)** Rotational averages of 3D projections of various γ-tubulin complex models (y-axis) using the indicated rotational fold-symmetries (x-axis). Averages are colored according to relative average intensity (arbitrary units). **E)** Scoring of the rotational averages in panel **D)** by power spectrum analysis. Each rotational average band (x-axis) is colored according to the average magnitude of the power spectrum, normalized such that for each structure, minimum values correspond to 0 and maximum values correspond to 1. γ-TuRC*_Ce_* (this study), human γ-TuRC in the “open” conformation (PDB ID: 6V6S (*20*)), and budding yeast γ-TuSC oligomers in the “closed” conformation (PDB ID: 5FLZ (*39*)) were analyzed. See Methods for further analysis details. **F)** Side and bottom views of the γ-TuRC*_Ce_* model rigid body fitted into the STA reconstruction of the minus end-capping structure in centrosomes of *C. elegans* mitotic embryos (EMD-19780) (*16*).

**Supplementary Figure 4.**
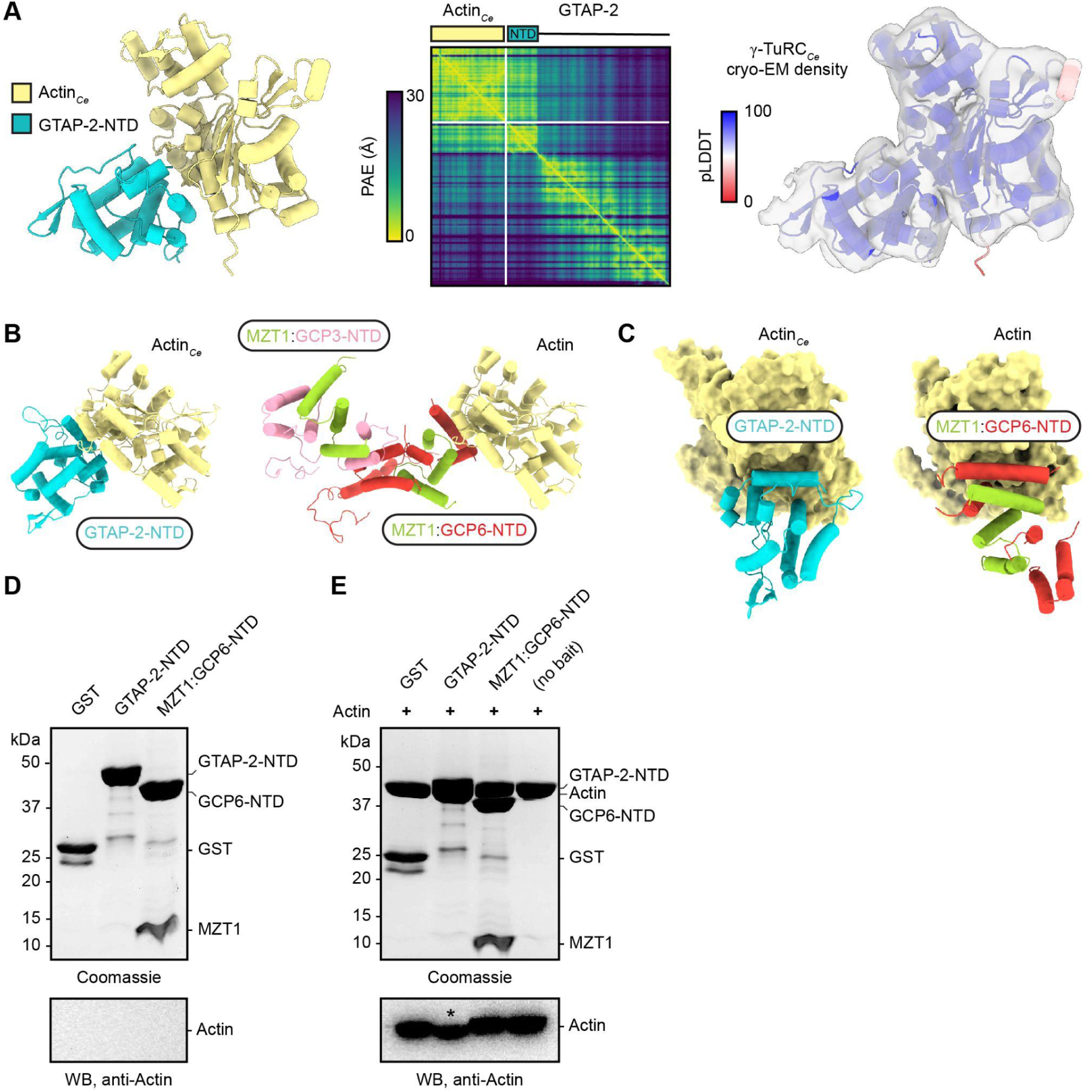
Details regarding the interaction between GTAP-2-NTD and actin*_Ce_*. **A)** Left: Cartoon representation of an AlphaFold3 prediction of full length of GTAP-2 with actin*_Ce_*. Only the GTAP-2-NTD is shown for clarity. Middle: the predicted aligned error (PAE) plot from the prediction. The region of the GTAP-2 sequence corresponding to GTAP-2-NTD is indicated in the schematic. Right: Cartoon representation the AlphaFold3 prediction colored according to the predicted local distance difference test (pLDDT) score and rigid body fitted into the γ-TuRC*_Ce_* cryo-EM density map (transparency surface; only cropped density is shown for clarity). Only the GTAP-2-NTD is shown for clarity. **B)** Left: Cartoon representation of actin*_Ce_* bound to GTAP-2-NTD from the γ-TuRC*_Ce_* model. Right: Cartoon representation of actin bound to MZT1:GCP3-NTD/MZT1:GCP6-NTD in the human γ-TuRC (PDB ID: 6X0U) (*21*). **C)** Left: View of GTAP-2-NTD (cartoon representation) bound to the barbed end groove of actin*_Ce_* (surface representation) from the γ-TuRC*_Ce_*model. Right: View of part of MZT1:GCP6-NTD (cartoon representation) bound to the barbed end groove of actin (surface representation) in the human γ-TuRC (PDB ID: 6X0U) (*21*). **D)** Coomassie-stained SDS-PAGE gel (top) of purified proteins used as baits in the GST pulldowns in Figure 2C. An anti-actin western blot (“WB”; bottom) confirms that actin is not present in the purified bait protein preparations. **E)** Coomassie-stained SDS-PAGE gel (top) and anti-actin western blot (bottom) of inputs of the GST pulldowns in Figure 2C. The actin band in the GTAP-2-NTD lane (indicated by an asterisk) is slightly lower than expected because it presumably overlaps with GST-tagged GTAP-2-NTD, which runs at a similar molecular weight as actin.

**Supplementary Figure 5.**
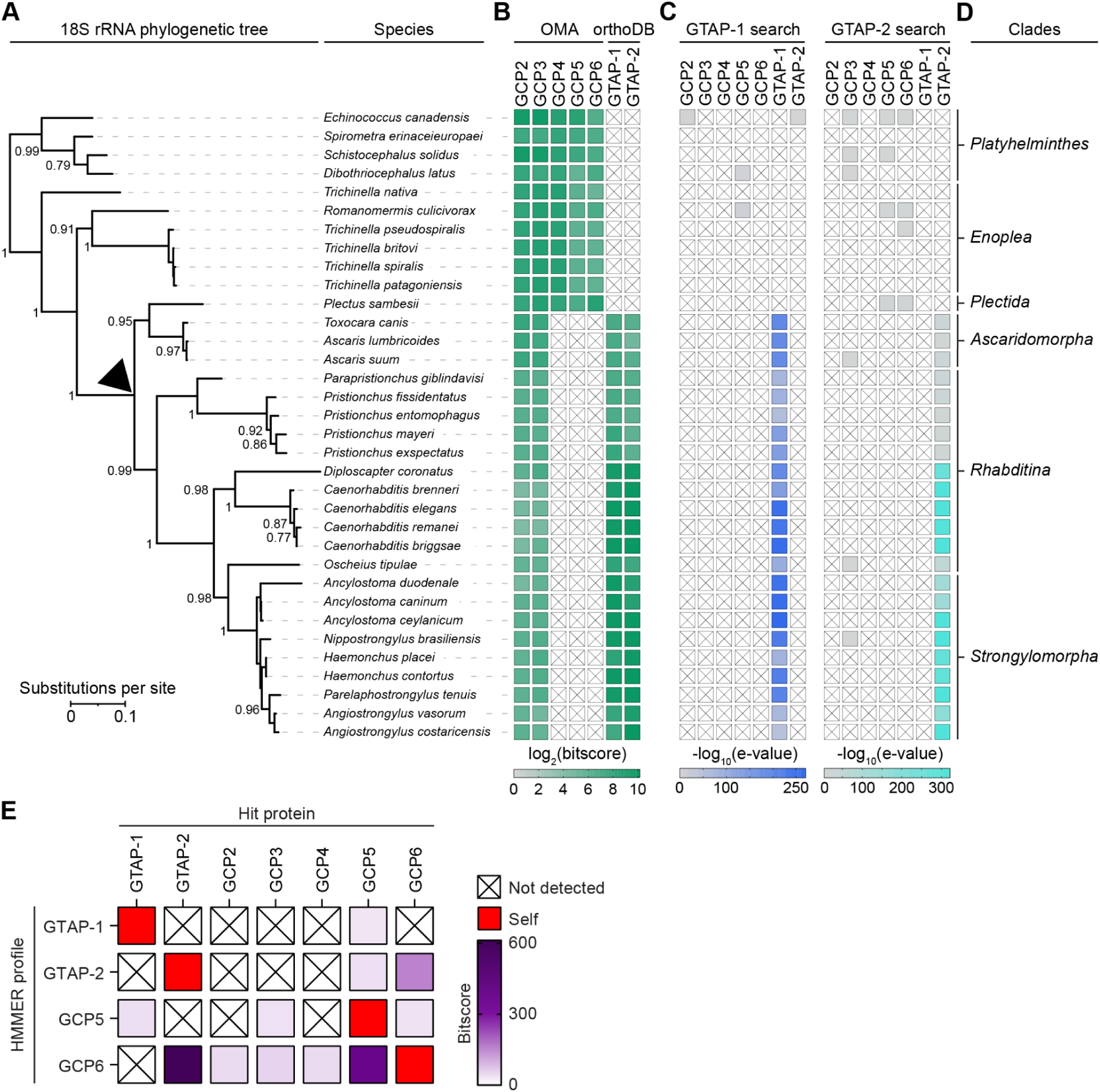
Phylogenetic evidence for the evolution of GTAP-1 and GTAP-2 from GCP5 and GCP6. **A)** Maximum likelihood-based phylogenetic tree constructed using RAxML (*80*) based on a multiple sequence alignment of 18S rRNA sequences. Node values indicate bootstrap support (fraction of 1,000 replicates). The scale bar represents 0.1 substitutions per site. The black triangle denotes the node closest to where a putative loss of the GCP4/5/4/6 assembly occurred. Tree visualization was performed using iTOL (*81*). **B)** Heatmap displaying protein bit scores for GCP2-6 (sourced from OMA; (*82*)) and GTAP-1-2 (sourced from OrthoDB; (*83*)) identified via protein-specific HMMER search (*84*). Color intensity scales with the bit score (1 to 1144). White cells marked with crosses indicate no ortholog detected. **C)** HMMER searches using GTAP-1 and GTAP-2 profile models demonstrate selective and specific detection of their corresponding orthologs across the analyzed species, with negligible cross-recognition of other GCP family members. **D)** Taxonomic clades of species from **A)**. **E)** Cross-HMMER validation matrix showing reciprocal specificity among GTAP-1, GTAP-2, GCP5, and GCP6 profile HMMs. Rows indicate the query HMMER profile used for the search, and respective hit proteins (taken from a single outgroup for GTAP-1/2 or GCP2/3/4/5/6) are shown in columns. Cells are colored by bitscore value. Red cells denote correct self-identification, whereas purple shading indicates cross-detection between related proteins. White cells marked with crosses indicate no significant hit detected.

**Supplementary Figure 6.**
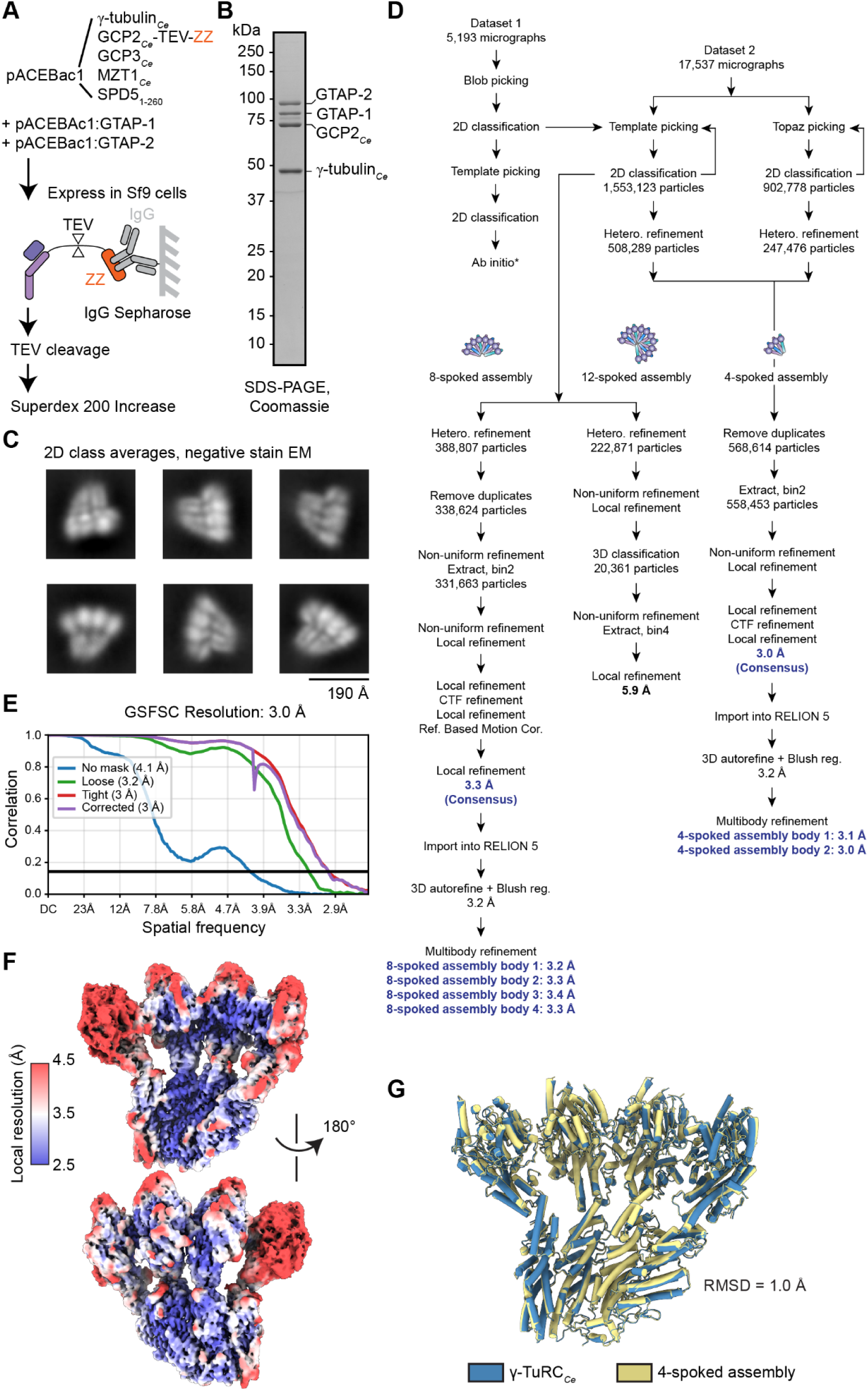
Purification and processing details for the 4-, 8-, and 12-spoked assembly cryo-EM reconstructions. **A)** Purification scheme for generating recombinant 4-spoked assemblies. **B)** Coomassie-stained SDS-PAGE gel showing peak gel filtration fraction corresponding to purified 4-spoked assemblies. Bands corresponding to 4-spoked assembly protein species are labeled. **C)** Six 2D class averages from negative stain EM of purified 4-spoked assemblies. **D)** SPA processing workflow for the 4-, 8-, and 12-spoked assemblies shown in Figures 3C and 5B. **E)** Gold standard FSC curve for the 4-spoked assembly cryo-EM reconstruction, generated in CryoSPARC (*70*). **F)** Frontside and backside views of the 4-spoked assembly cryo-EM reconstruction colored according to local resolution, calculated in CryoSPARC (*70*). **G)** Aligned models of the 4-spoked assembly from the ã-TuRC*_Ce_* model (blue) or from the reconstituted subcomplex in Figure 3C (yellow), shown in cartoon representation. The RMSD from the alignment as calculated by ChimeraX is indicated (*74*).

**Supplementary Figure 7.**
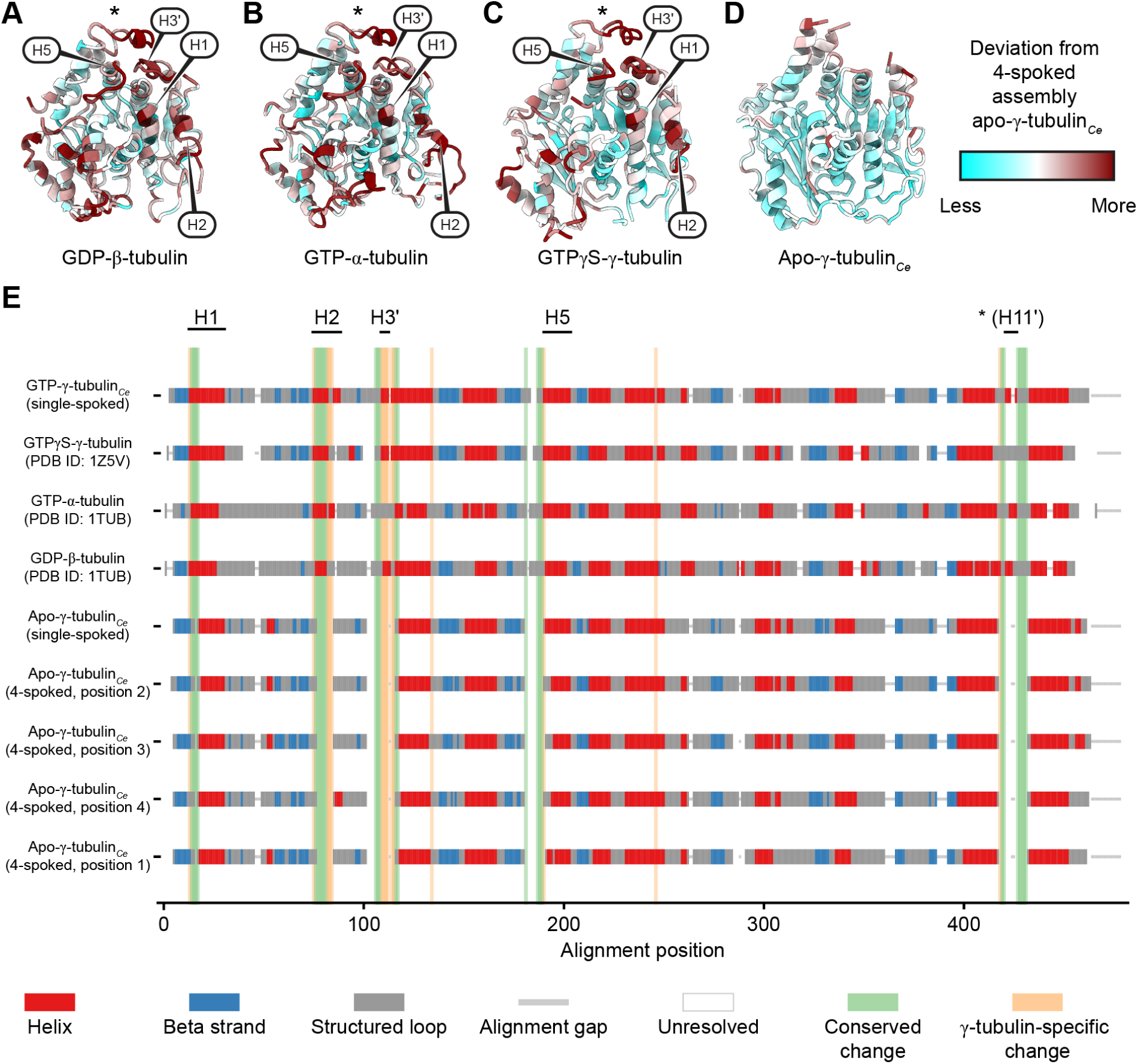
Comparison of apo-γ-tubulin*_Ce_* with nucleotide-bound α-, β- and γ-tubulin structures. **A)**-**D)** Cartoon representations of GDP-β-tubulin (**A**; PDB ID: 1TUB (*46*)), GTP-α-tubulin (**B**; PDB ID: 1TUB (*46*)), GTPγS-γ-tubulin (**C**; PDB ID: 1Z5V (*27*)), and apo-γ-tubulin*_Ce_* (**D**), from the 4-spoked assembly density map (Figure 4A), as a control. Models are colored according to the output of the “measure mapvalues” command in ChimeraX, as in the right of panel **D)** and in Figure 4B. Locations of helices H1, H2, H3’, and H5 are indicated in panels **A)**-**C)**. **E)** Secondary structure vs. primary sequence for the various α-, β- and γ-tubulin structures examined in this study. Secondary structure features were defined by the Define Secondary Structure of Proteins (DSSP) algorithm via the Biopython structural bioinformatics module (*85*, *86*). Tubulin primary sequence alignments were pre-computed with the MAFFT E-INS-i algorithm (*87*), and structural features returned by DSSP were imposed onto the alignment. Unresolved residues lacking coordinates in the 3D structures were flagged as structural gaps, while sequence alignment gaps were preserved and are displayed. Automated position-by-position comparative logic was applied to detect structural transitions in the nucleotide-bound complexes against a baseline established by the five apo-γ-tubulin*_Ce_* models derived from the reconstituted single or 4-spoked assemblies. Conserved changes (green) represent secondary structure differences between all compared structures, while γ-tubulin-specific changes (orange) represent only those differences between γ-tubulin models (including PDB ID: 1Z5V (*27*)).

**Supplementary Figure 8.**
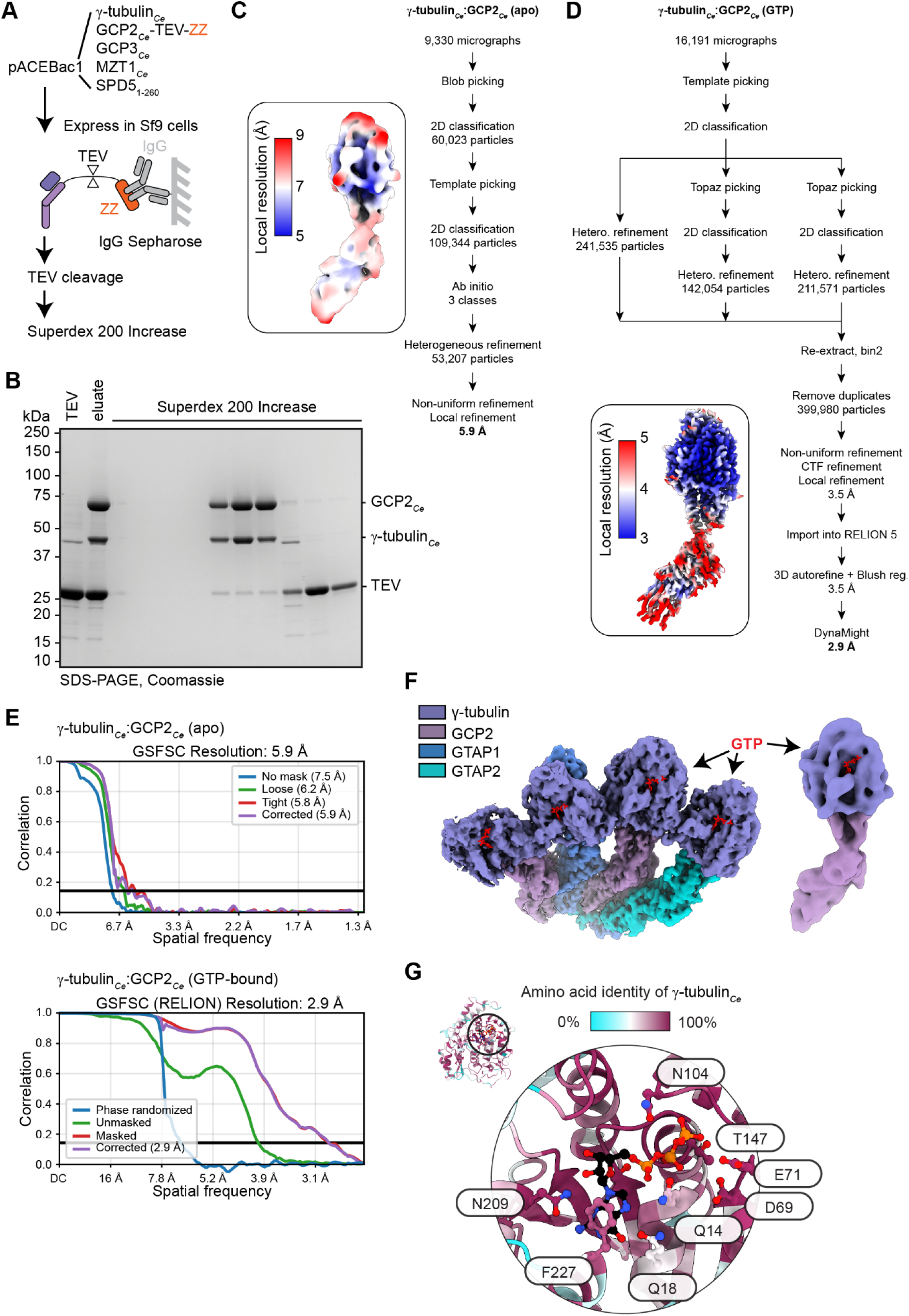
Purification and processing details for γ-tubulin*_Ce_*:GCP2*_Ce_*cryo-EM reconstructions. **A)** Purification scheme for generating recombinant γ-tubulin*_Ce_*:GCP2*_Ce_* single-spoked assemblies. **B)** Coomassie-stained SDS-PAGE gel showing gel filtration fractions of γ-tubulin*_Ce_*:GCP2*_Ce_*. Bands corresponding to γ-tubulin*_Ce_*, GCP2*_Ce_* and tobacco etch virus (TEV) protease are labeled. **C)-D)** SPA processing workflow for the apo (**C**) and GTP-bound (**D**) γ-tubulin*_Ce_*:GCP2*_Ce_*reconstructions. Insets show the respective cryo-EM reconstructions colored according to local resolution, calculated using CryoSPARC (*70*). **E)** Gold standard FSC curve for the apo (top, generated using CryoSPARC (*70*)) and GTP-bound (bottom, calculated using RELION 5 (*76*)) γ-tubulin*_Ce_*:GCP2*_Ce_* reconstructions. **F)** Nucleotide binding site views of the recombinant 4-spoked assembly reconstruction (left) and the apo γ-tubulin*_Ce_*:GCP2*_Ce_* reconstruction (right) colored according to the legend. GTP is shown in red stick representation at the expected location to highlight the obvious lack of density for nucleotide. **G)** Zoomed-in view of γ-tubulin*_Ce_*’s nucleotide binding site, colored according to conservation evaluated against a multiple sequence alignment of 2,000 γ-tubulin sequences obtained from a random selection of species from UniProt (*88*). Conserved residues involved in coordinating the guanine nucleotide are labeled and shown in ball and stick representation and colored by heteroatom. GTP is shown in black ball and stick representation, additionally colored by heteroatom.

**Supplementary Figure 9.**
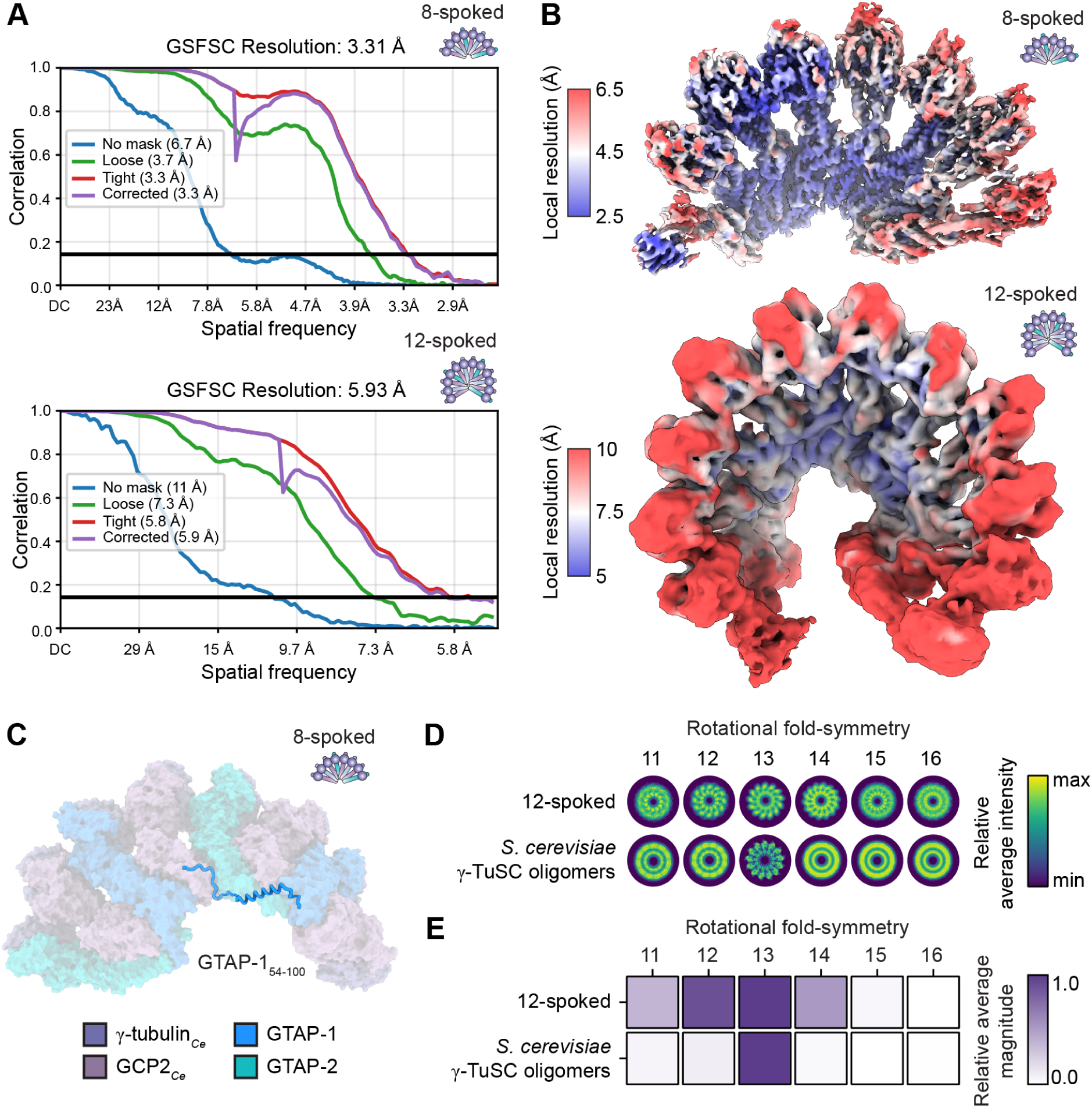
Resolution and architecture analysis of the 8- and 12-spoked assembly cryo-EM reconstructions and models. **A)** Gold standard FSC curve for 8-spoked (top) and 12-spoked (bottom) assembly cryo-EM reconstructions, generated in CryoSPARC (*70*). **B)** Top views of the 8-spoked (top) and 12-spoked (bottom) assembly cryo-EM reconstructions colored according to local resolution, as calculated in CryoSPARC (*70*). **C)** The 8-spoked assembly model (faded surface representation, schematized in the top right corner) viewed from the bottom, with the extension of the GTAP-1 N-terminus (residues 54-100) shown in cartoon representation. Subunits are colored according to the legend. **D)** Rotational averages of 3D projections of various γ-tubulin complex models (y-axis) using the indicated rotational fold-symmetries (x-axis). Averages are colored according to relative average intensity (arbitrary units). **E)** Scoring of the rotational averages in panel **D)** by power spectrum analysis. Each rotational average band (x-axis) is colored according to the average magnitude of the power spectrum, normalized such that for each structure, minimum values correspond to 0 and maximum values correspond to 1. The 12-spoked assembly (this study) or budding yeast γ-TuSC oligomers in the “closed” conformation (PDB ID: 5FLZ (*39*)) were analyzed. See Methods for further analysis details.

**Supplementary Figure 10.**
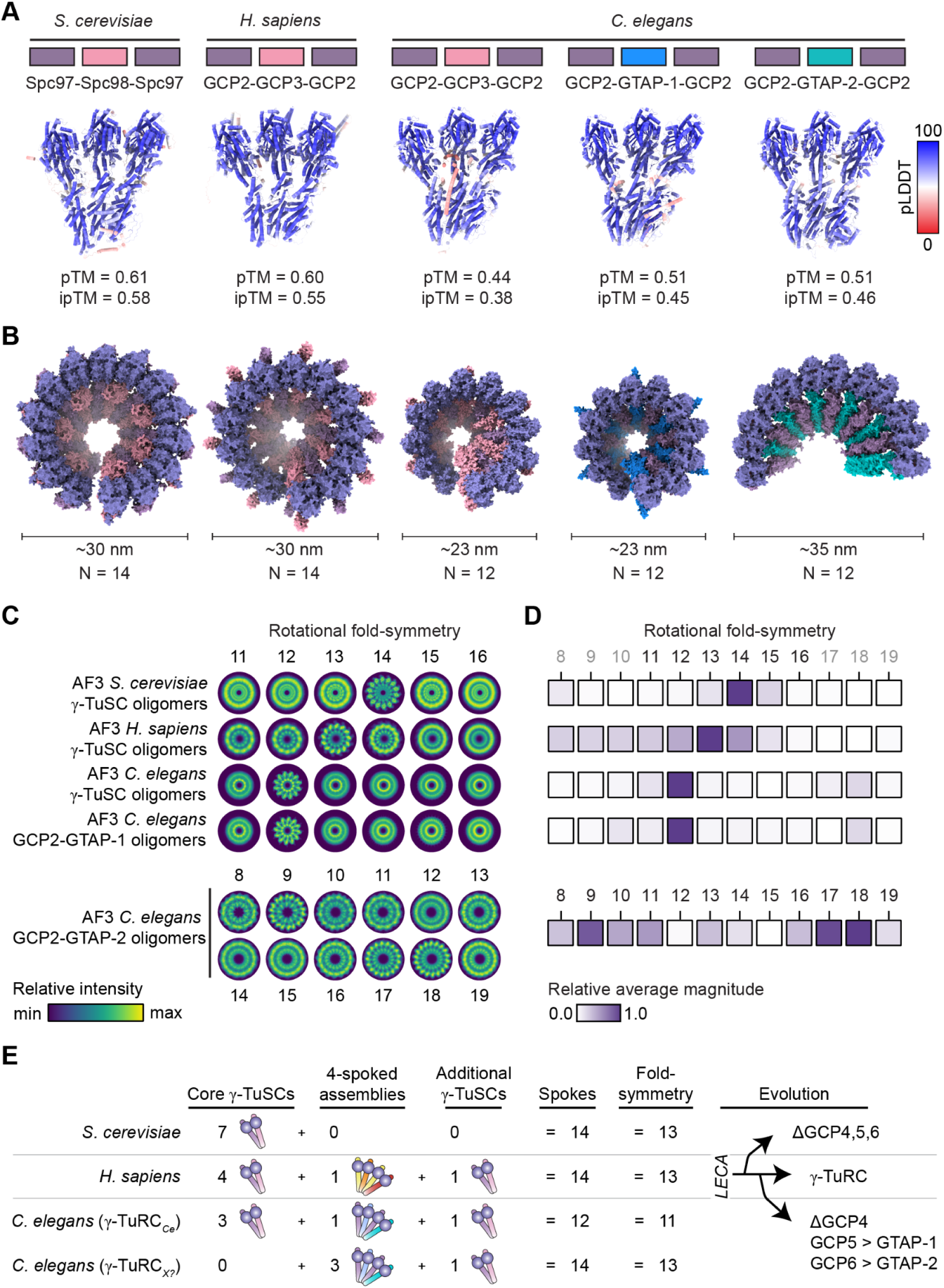
Consequences and implications from architectural switching in the evolution of C. elegans γ-tubulin complexes. **A)** AlphaFold 3 predictions containing three copies of γ-tubulin, two copies of GCP2, and one copy of GCP3, GTAP-1 or GTAP-2 from *S. cerevisiae* (left), *H. sapiens* (middle), and *C. elegans* (right) (*41*). Models are shown in cartoon representation and colored according to pLDDT. **B)** *In silico* γ-tubulin complexes obtained by propagation of models in **A).** Neighboring models were aligned by the first and last GCP2 to generate *in silico* γ-TuRC-like assemblies containing the indicated number of spokes (N). Approximate diameter of resulting assemblies is indicated. **C)** Rotational averages of 3D projections of *in silico* γ-tubulin complex models from **B)** (y-axis) using the indicated rotational fold-symmetries (x-axis). Averages are colored according to relative average intensity (arbitrary units). **D)** Scoring of the rotational averages in panel **C)** by power spectrum analysis. Each rotational average band (x-axis) is colored according to the average magnitude of the power spectrum, normalized such that for each structure, minimum values correspond to 0 and maximum values correspond to 1. γ-TuRC*_Ce_* (this study), human γ-TuRC (PDB ID: 6V6S (*20*)), and budding yeast γ-TuSC oligomers (PDB ID: 5FLZ (*39*)) were analyzed. See Methods for further analysis details. **E)** An architectural switch in the evolution of the ã-TuRC. Schematic comparison of ã-TuRC composition in *S. cerevisiae*, *H. sapiens*, and *C. elegans*. In *S. cerevisiae*, seven γ-TuSCs assemble into a 14-spoked γ-tubulin ring that provides a 13-protofilament nucleation template (*42*). In humans, a total of five γ-TuSCs and a single 4-spoked GCP4/5/4/6 assembly form the canonical γ-TuRC, also yielding a 14-spoked ring with 13-fold radial symmetry (*20*). In *C. elegans*, the dominant γ-TuRC species (γ-TuRC*_Ce_*) is most likely assembled from a total of four γ-TuSCs surrounding an evolutionary divergent 4-spoke module containing GCP2*_Ce_*, GTAP-1 and GTAP-2, instead of GCP4/5/4/6. This results in an architecturally distinct, ∼11-protofilament nucleation template built from a 12-spoked assembly. In addition to γ-TuRC*_Ce_*, however, evolution in the *C. elegans* GCPs uniquely allows for other assemblies to theoretically form (using e.g. multiple 4-spoked oligomers), which can switch their architectures back to accommodating ∼13-protofilament microtubule lattices (γ-TuRC*_X_*). LECA: Last Eukaryotic Common Ancestor.

## Supplementary Tables

**Supplementary Table 1.** Mass spectrometry analysis of γ-TuRC*_Ce_*.

| Subunit | Accession reported by LC-MS/MS | Unique peptides | Coverage | iBAQ ratio relative to 6 x GCP2<sub>Ce</sub> |
| --- | --- | --- | --- | --- |
| γ-tubulin<sub>Ce</sub> | P34475 | 17 | 55% | 19.0 |
| GCP2<sub>Ce</sub> | G5EF84 | 27 | 57% | 6 |
| GCP3<sub>Ce</sub> | O61208 | 21 | 26% | 1.1 |
| GTAP-1 | n.d.* | 17 | 24% | 1.2 |
| GTAP-2 | P34651 | 27 | 40% | 1.8 |
| Actin<sub>Ce</sub> | P0DM41 | 14 | 39% | 2.9 |
| MZT1<sub>Ce</sub> | n.d. | n.d. | n.d. | n.d. |
| SPD-5 | n.d. | n.d. | n.d. | n.d. |

\*Accession code available at UniProt: Q23421; n.d. = not detected

**Supplementary Table 2.** γ-TuRC*_Ce_* cryo-EM data collection and processing statistics.

|  |  |
| --- | --- |
| Magnification | 130,000 X |
| Voltage (keV) | 300 |
| Electron exposure (e $\text{\AA}^{-2}$) | 40 |
| Defocus range ($\mu\text{m}$) | -0.8 to -2.6 |
| Pixel size ($\text{\AA}$) | 0.65 |
| Camera | Gatan K3 |
| Symmetry imposed | C1 |
| GIF slit width | -20 eV |
| No. initial particle images | 144,561 |
| No. final particle images | 21,493 |
| Map resolution ($\text{FSC}_{0.143}$; $\text{\AA}$) | 6.1 |
| EMDB ID | 58822 |

**Supplementary Table 3.** γ-TuRC*_Ce_* model building statistics.

|  |  |
| --- | --- |
| Initial model used | AlphaFold and 4-spoked assembly |
| Model resolution (FSC<sub>0.143</sub>; Å) | 7.3 |
| Map sharpening *B* factor (Å<sup>2</sup>) | 0 |
| **Model composition** |  |
| Non-hydrogen atoms | 77,125 |
| Protein residues | 9,620 |
| Ligands | 0 |
| **B factors (Å<sup>2</sup>)** |  |
| Protein | 607.8 |
| Ligand | n/a |
| **R.m.s. deviations from ideality** |  |
| Bonds (Å) | 0.003 |
| Angles (°) | 0.672 |
| *MolProbity* score | 2.20 |
| Clashscore | 22.04 |
| Poor rotamers (%) | 0.02 |
| **Ramachandran plot** |  |
| Favoured (%) | 94.75 |
| Allowed (%) | 5.18 |
| Disallowed (%) | 0.07 |
| PDB ID | 32DT |

**Supplementary Table 4.** 4-spoked assembly cryo-EM data collection and processing statistics.

|  |  |
| --- | --- |
| Magnification | 130,000 X |
| Voltage (keV) | 300 |
| Electron exposure (e Å<sup>-2</sup>) | 31 |
| Defocus range (μm) | -0.8 to -2.6 |
| Pixel size (Å) | 0.65 |
| Camera | Gatan K3 |
| Symmetry imposed | C1 |
| GIF slit width | -20 eV |
| No. initial particle images | 2,455,901 (4-spoked assembly)<br>1,553,123 (8- and 12-spoked assemblies) |
| No. final particle images | 558,431 (4-spoked assembly);<br>330,106 (8-spoked assembly);<br>20,361 (12-spoked assembly) |
| Map resolution (FSC<sub>0.143</sub>, Å) | 3.0 (4-spoked assembly);<br>3.3 (8-spoked assembly);<br>5.9 (12-spoked assembly) |
| EMDB ID | 58888 (4-spoked assembly);<br>58884 (8-spoked assembly);<br>58828 (12-spoked assembly) |

**Supplementary Table 5.** 4- and 8-spoked assembly model building statistics.

|  | **4-spoked assembly** | **8-spoked assembly** |
| --- | --- | --- |
| Initial model used | AlphaFold | 4-spoked assembly |
| Model resolution (FSC<sub>0.143</sub>; Å) | 3.0 | 3.3 |
| Map sharpening *B* factor (Å<sup>2</sup>) | 0 | 0 |
| **Model composition** |  |  |
| Non-hydrogen atoms | 31,777 | 63,679 |
| Protein residues | 3,932 | 7,879 |
| Ligands | 0 | 0 |
| **B factors (Å<sup>2</sup>)** |  |  |
| Protein | 112.22 | 173.16 |
| Ligand | n/a | n/a |
| **R.m.s. deviations from ideality** |  |  |
| Bonds (Å) | 0.010 | 0.004 |
| Angles (°) | 0.609 | 0.625 |
| *MolProbity* score | 1.98 | 1.99 |
| Clashscore | 6.93 | 9.59 |
| Poor rotamers (%) | 2.85 | 1.95 |
| **Ramachandran plot** |  |  |
| Favoured (%) | 96.28 | 96.16 |
| Allowed (%) | 3.72 | 3.79 |
| Disallowed (%) | 0.00 | 0.05 |
| PDB ID | 32HE | 32GX |

**Supplementary Table 6.** γ-tubulin*_Ce_*:GCP2*_Ce_* cryo-EM data collection and processing statistics.

|  | **Apo** | **GTP** |
| --- | --- | --- |
| Magnification | 130,000 X | 130,000 X |
| Voltage (keV) | 300 | 300 |
| Electron exposure (e $\text{\AA}^{-2}$) | 59 | 30 |
| Defocus range ($\mu\text{m}$) | -0.5 to -2.5 | -0.6 to -2.6 |
| Pixel size ($\text{\AA}$) | 0.65 | 0.65 |
| Camera | Gatan K3 | Gatan K3 |
| Symmetry imposed | C1 | C1 |
| GIF slit width | -20 eV | -20 eV |
| No. initial particle images | 308,217 | 1,689,660 |
| No. final particle images | 53,207 | 399,980 |
| Map resolution ($\text{FSC}_{0.143}$, $\text{\AA}$) | 5.9 | 2.9 |
| EMDB ID | 58833 | 58866 |

**Supplementary Table 7.**
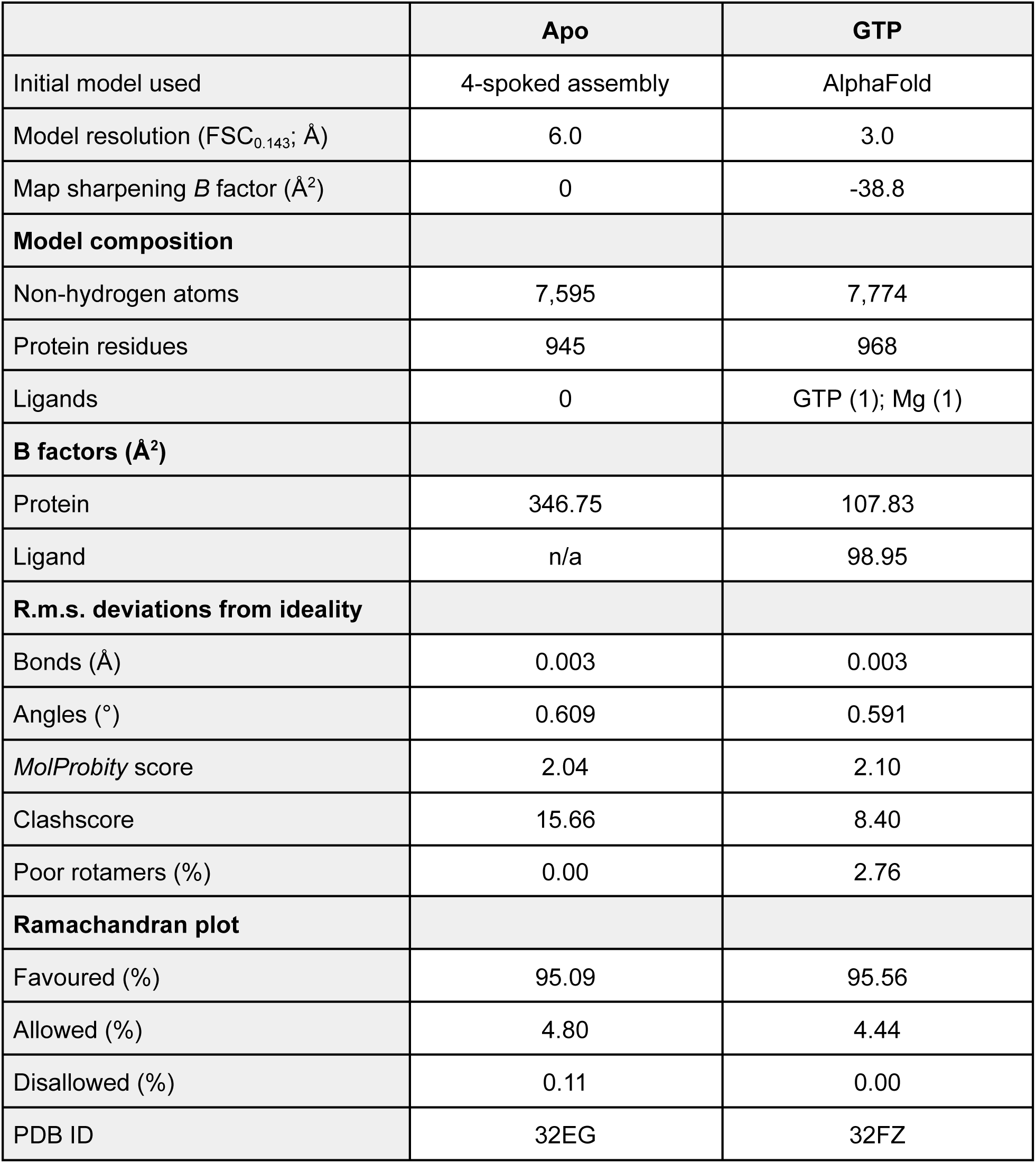
γ-tubulin*_Ce_*:GCP2*_Ce_* model building statistics.

|  | **Apo** | **GTP** |
| --- | --- | --- |
| Initial model used | 4-spoked assembly | AlphaFold |
| Model resolution (FSC<sub>0.143</sub>; Å) | 6.0 | 3.0 |
| Map sharpening *B* factor (Å<sup>2</sup>) | 0 | -38.8 |
| **Model composition** |  |  |
| Non-hydrogen atoms | 7,595 | 7,774 |
| Protein residues | 945 | 968 |
| Ligands | 0 | GTP (1); Mg (1) |
| **B factors (Å<sup>2</sup>)** |  |  |
| Protein | 346.75 | 107.83 |
| Ligand | n/a | 98.95 |
| **R.m.s. deviations from ideality** |  |  |
| Bonds (Å) | 0.003 | 0.003 |
| Angles (°) | 0.609 | 0.591 |
| *MolProbity* score | 2.04 | 2.10 |
| Clashscore | 15.66 | 8.40 |
| Poor rotamers (%) | 0.00 | 2.76 |
| **Ramachandran plot** |  |  |
| Favoured (%) | 95.09 | 95.56 |
| Allowed (%) | 4.80 | 4.44 |
| Disallowed (%) | 0.11 | 0.00 |
| PDB ID | 32EG | 32FZ |

## Other Supplementary Information

**Supplementary Video 1. Comparison between apo γ-tubulin*_Ce_*from the 4-spoked assembly reconstruction with γ-tubulin*_Ce_* from the GTP-γ-tubulin*_Ce_*:GCP2*_Ce_* reconstruction.** The movie shows the apo γ-tubulin*_Ce_* model (cartoon representation) first, then the corresponding density, then GTP (red ball and stick representation) from the GTP-γ-tubulin*_Ce_*:GCP2*_Ce_* model only. The apo γ-tubulin*_Ce_* density then morphs into the GTP-γ-tubulin*_Ce_* density, followed by the GTP-γ-tubulin*_Ce_* model. The last frames of the movie show the difference map between zoned the apo γ-tubulin*_Ce_* and the GTP-γ-tubulin*_Ce_* densities (maroon surfaces) overlaid on the GTP-γ-tubulin*_Ce_* model to highlight the regions that change between the two structures.

## References

1. Y. Zheng, M. L. Wong, B. Alberts, T. Mitchison, Nucleation of microtubule assembly by a gamma-tubulin-containing ring complex. Nature 378, 578–583 (1995).

2. T. J. Keating, G. G. Borisy, Immunostructural evidence for the template mechanism of microtubule nucleation. Nat. Cell Biol. 2, 352–357 (2000).

3. A. Aher, L. Urnavicius, A. Xue, K. Neselu, T. M. Kapoor, Structure of the γ-tubulin ring complex-capped microtubule. Nat. Struct. Mol. Biol., doi: 10.1038/s41594-024-01264-z (2024).

4. M. C. Ledbetter, K. R. Porter, A “microtubule” in plant cell fine structure. J. Cell Biol. 19, 239–250 (1963).

5. L. G. Tilney, J. Bryan, D. J. Bush, K. Fujiwara, M. S. Mooseker, D. B. Murphy, D. H. Snyder, Microtubules: evidence for 13 protofilaments. J. Cell Biol. 59, 267–275 (1973).

6. E. Unger, K. J. Böhm, W. Vater, Structural diversity and dynamics of microtubules and polymorphic tubulin assemblies. Electron Microsc. Rev. 3, 355–395 (1990).

7. S. Chaaban, G. J. Brouhard, A microtubule bestiary: structural diversity in tubulin polymers. Mol. Biol. Cell 28, 2924–2931 (2017).

8. J. L. Ferreira, V. Pražák, D. Vasishtan, M. Siggel, F. Hentzschel, A. M. Binder, E. Pietsch, J. Kosinski, F. Frischknecht, T. W. Gilberger, K. Grünewald, Variable microtubule architecture in the malaria parasite. Nat. Commun. 14, 1216 (2023).

9. P. R. Burton, R. Hinkley, G. Pierson, Tannic acid-stained microtubules with 12, 13, and 15 protofilaments. J. Cell Biol. 65, 227–233 (1975).

10. Z. Xu, B. A. Afzelius, Early changes in the substructure of the marginal bundle in human blood platelets responding to adenosine diphosphate. Journal of ultrastructure and molecular structure research 99, 254–260 (1988).

11. C. Savage, M. Hamelin, J. G. Culotti, A. Coulson, D. G. Albertson, M. Chalfie, mec-7 is a beta-tubulin gene required for the production of 15-protofilament microtubules in Caenorhabditis elegans. Genes Dev. 3, 870–881 (1989).

12. E. C. Raff, J. D. Fackenthal, J. A. Hutchens, H. D. Hoyle, F. R. Turner, Microtubule architecture specified by a beta-tubulin isoform. Science 275, 70–73 (1997).

13. S.-C. Ti, G. M. Alushin, T. M. Kapoor, Human β-Tubulin Isotypes Can Regulate Microtubule Protofilament Number and Stability. Dev. Cell 47, 175–190.e5 (2018).

14. J. G. Cueva, J. Hsin, K. C. Huang, M. B. Goodman, Posttranslational acetylation of α-tubulin constrains protofilament number in native microtubules. Curr. Biol. 22, 1066–1074 (2012).

15. M. Chalfie, J. N. Thomson, Structural and functional diversity in the neuronal microtubules of Caenorhabditis elegans. J. Cell Biol. 93, 15–23 (1982).

16. F. Tollervey, M. U. Rios, E. Zagoriy, J. B. Woodruff, J. Mahamid, Molecular architectures of centrosomes in C. elegans embryos visualized by cryo-electron tomography. Dev. Cell 60, 885–900.e5 (2025).

17. S. Chaaban, S. Jariwala, C.-T. Hsu, S. Redemann, J. M. Kollman, T. Müller-Reichert, D. Sept, K. H. Bui, G. J. Brouhard, The Structure and Dynamics of C. elegans Tubulin Reveals the Mechanistic Basis of Microtubule Growth. Dev. Cell 47, 191–204.e8 (2018).

18. M. Chalfie, J. N. Thomson, Organization of neuronal microtubules in the nematode Caenorhabditis elegans. J. Cell Biol. 82, 278–289 (1979).

19. T. Fukushige, Z. K. Siddiqui, M. Chou, J. G. Culotti, C. B. Gogonea, S. S. Siddiqui, M. Hamelin, MEC-12, an alpha-tubulin required for touch sensitivity in C. elegans. J. Cell Sci. 112 (Pt 3), 395–403 (1999).

20. M. Wieczorek, L. Urnavicius, S.-C. Ti, K. R. Molloy, B. T. Chait, T. M. Kapoor, Asymmetric Molecular Architecture of the Human γ-Tubulin Ring Complex. Cell, doi: 10.1016/j.cell.2019.12.007 (2019).

21. M. Wieczorek, T.-L. Huang, L. Urnavicius, K.-C. Hsia, T. M. Kapoor, MZT Proteins Form Multi-Faceted Structural Modules in the γ-Tubulin Ring Complex. Cell Rep. 31, 107791 (2020).

22. P. Liu, E. Zupa, A. Neuner, A. Böhler, J. Loerke, D. Flemming, T. Ruppert, T. Rudack, C. Peter, C. Spahn, O. J. Gruss, S. Pfeffer, E. Schiebel, Insights into the assembly and activation of the microtubule nucleator γ-TuRC. Nature 578, 467–471 (2019).

23. T. Consolati, J. Locke, J. Roostalu, Z. A. Chen, J. Gannon, J. Asthana, W. M. Lim, F. Martino, M. A. Cvetkovic, J. Rappsilber, A. Costa, T. Surrey, Microtubule Nucleation Properties of Single Human γTuRCs Explained by Their Cryo-EM Structure. Dev. Cell 53, 603–617.e8 (2020).

24. F. Zimmermann, M. Serna, A. Ezquerra, R. Fernandez-Leiro, O. Llorca, J. Luders, Assembly of the asymmetric human γ-tubulin ring complex by RUVBL1-RUVBL2 AAA ATPase. Sci Adv 6 (2020).

25. K. Oegema, C. Wiese, O. C. Martin, R. A. Milligan, A. Iwamatsu, T. J. Mitchison, Y. Zheng, Characterization of two related Drosophila gamma-tubulin complexes that differ in their ability to nucleate microtubules. J. Cell Biol. 144, 721–733 (1999).

26. L. M. Rice, E. A. Montabana, D. A. Agard, The lattice as allosteric effector: structural studies of alphabeta- and gamma-tubulin clarify the role of GTP in microtubule assembly. Proc. Natl. Acad. Sci. U. S. A. 105, 5378–5383 (2008).

27. H. Aldaz, L. M. Rice, T. Stearns, D. A. Agard, Insights into microtubule nucleation from the crystal structure of human gamma-tubulin. Nature 435, 523–527 (2005).

28. M. Würtz, E. Zupa, E. S. Atorino, A. Neuner, A. Böhler, A. S. Rahadian, B. J. A. Vermeulen, G. Tonon, S. Eustermann, E. Schiebel, S. Pfeffer, Modular assembly of the principal microtubule nucleator γ-TuRC. Nat. Commun. 13, 473 (2022).

29. E. Hannak, K. Oegema, M. Kirkham, P. Gönczy, B. Habermann, A. A. Hyman, The kinetically dominant assembly pathway for centrosomal asters in Caenorhabditis elegans is gamma-tubulin dependent. J. Cell Biol. 157, 591–602 (2002).

30. M. D. Sallee, J. C. Zonka, T. D. Skokan, B. C. Raftrey, J. L. Feldman, Tissue-specific degradation of essential centrosome components reveals distinct microtubule populations at microtubule organizing centers. PLoS Biol. 16, e2005189 (2018).

31. N. Haruta, E. Sumiyoshi, Y. Honda, M. Terasawa, C. Uchiyama, M. Toya, Y. Kubota, A. Sugimoto, A germline-specific role for unconventional components of the γ-tubulin complex in Caenorhabditis elegans. J. Cell Sci. 136 (2023).

32. M. Ohta, Z. Zhao, D. Wu, S. Wang, J. L. Harrison, J. S. Gómez-Cavazos, A. Desai, K. F. Oegema, Polo-like kinase 1 independently controls microtubule-nucleating capacity and size of the centrosome. J. Cell Biol. 220 (2021).

33. M. Ohta, O. Arakawa, Y. Gu, W. Tian, K. D. Corbett, A. Desai, K. Oegema, Phosphorylation remodels the mitotic centrosome matrix to generate bipartite γ-tubulin complex docking sites, bioRxivorg (2025). 10.1101/2025.11.20.689565.

34. R. A. Walker, E. T. O’brien, N. K. Pryer, M. F. Soboeiro, W. A. Voter, H. P. Erickson, E. D. Salmon, Dynamic instability of individual microtubules analyzed by video light microscopy: rate constants and transition frequencies. J. Cell Biol. 107, 1437–1448 (1988).

35. M. Wieczorek, S.-C. Ti, L. Urnavicius, K. R. Molloy, A. Aher, B. T. Chait, T. M. Kapoor, Biochemical reconstitutions reveal principles of human γ-TuRC assembly and function. J. Cell Biol. 220 (2021).

36. Y. Xu, H. Muñoz-Hernández, R. Krutyhołowa, F. Marxer, F. Cetin, M. Wieczorek, Partial closure of the γ-tubulin ring complex by CDK5RAP2 activates microtubule nucleation. Dev. Cell 0 (2024).

37. A. Thawani, M. J. Rale, N. Coudray, G. Bhabha, H. A. Stone, J. W. Shaevitz, S. Petry, The transition state and regulation of γ-TuRC-mediated microtubule nucleation revealed by single molecule microscopy. Elife 9 (2020).

38. H. Sui, K. H. Downing, Structural basis of interprotofilament interaction and lateral deformation of microtubules. Structure 18, 1022–1031 (2010).

39. J. M. Kollman, C. H. Greenberg, S. Li, M. Moritz, A. Zelter, K. K. Fong, J.-J. Fernandez, A. Sali, J. Kilmartin, T. N. Davis, D. A. Agard, Ring closure activates yeast γTuRC for species-specific microtubule nucleation. Nat. Struct. Mol. Biol. 22, 132–137 (2015).

40. F.-Q. Zhao, R. Craig, Capturing time-resolved changes in molecular structure by negative staining. J. Struct. Biol. 141, 43–52 (2003).

41. J. Abramson, J. Adler, J. Dunger, R. Evans, T. Green, A. Pritzel, O. Ronneberger, L. Willmore, A. J. Ballard, J. Bambrick, S. W. Bodenstein, D. A. Evans, C.-C. Hung, M. O’Neill, D. Reiman, K. Tunyasuvunakool, Z. Wu, A. Žemgulytė, E. Arvaniti, C. Beattie, O. Bertolli, A. Bridgland, A. Cherepanov, M. Congreve, A. I. Cowen-Rivers, A. Cowie, M. Figurnov, F. B. Fuchs, H. Gladman, R. Jain, Y. A. Khan, C. M. R. Low, K. Perlin, A. Potapenko, P. Savy, S. Singh, A. Stecula, A. Thillaisundaram, C. Tong, S. Yakneen, E. D. Zhong, M. Zielinski, A. Žídek, V. Bapst, P. Kohli, M. Jaderberg, D. Hassabis, J. M. Jumper, Accurate structure prediction of biomolecular interactions with AlphaFold 3. Nature 630, 493–500 (2024).

42. J. M. Kollman, J. K. Polka, A. Zelter, T. N. Davis, D. A. Agard, Microtubule nucleating gamma-TuSC assembles structures with 13-fold microtubule-like symmetry. Nature 466, 879–882 (2010).

43. A. Zheng, B. J. A. Vermeulen, M. Würtz, A. Neuner, N. Lübbehusen, M. P. Mayer, E. Schiebel, S. Pfeffer, Structural insights into the interplay between microtubule polymerases, γ-tubulin complexes and their receptors. Nat. Commun. 16, 402 (2025).

44. J. He, T. Li, S.-Y. Huang, Improvement of cryo-EM maps by simultaneous local and non-local deep learning. Nat. Commun. 14, 3217 (2023).

45. D. B. N. Vinh, J. W. Kern, W. O. Hancock, J. Howard, T. N. Davis, Reconstitution and characterization of budding yeast gamma-tubulin complex. Mol. Biol. Cell 13, 1144–1157 (2002).

46. E. Nogales, S. G. Wolf, K. H. Downing, Structure of the alpha beta tubulin dimer by electron crystallography. Nature 391, 199–203 (1998).

47. J. Schwab, D. Kimanius, A. Burt, T. Dendooven, S. H. W. Scheres, DynaMight: estimating molecular motions with improved reconstruction from cryo-EM images. Nat. Methods 21, 1855–1862 (2024).

48. C. Yin, W. Tang, Y. Fu, R. Z. Qi, FRET-based analysis of guanine nucleotide binding to γ-tubulin. Mol. Biol. Cell 37, ar46 (2026).

49. A. S. Kennard, K. B. Velle, R. Ranjan, D. Schulz, L. K. Fritz-Laylin, Tubulin sequence divergence is associated with the use of distinct microtubule regulators. Curr. Biol. 35, 233–248.e8 (2025).

50. M. Serna, F. Zimmermann, C. Vineethakumari, N. Gonzalez-Rodriguez, O. Llorca, J. Lüders, CDK5RAP2 activates microtubule nucleator γTuRC by facilitating template formation and actin release. Dev. Cell 0 (2024).

51. H. Muñoz-Hernández, Y. Xu, A. Pellicer Camardiel, D. Zhang, A. Xue, A. Aher, E. Walker, F. Marxer, T. M. Kapoor, M. Wieczorek, Structure of the microtubule-anchoring factor NEDD1 bound to the γ-tubulin ring complex. J. Cell Biol. 224 (2025).

52. M. Wieczorek, Conformational regulation of vertebrate γ-tubulin ring complexes by CM1 proteins. Cytoskeleton (Hoboken) 82, 513–515 (2025).

53. C. A. Tovey, C. E. Tubman, E. Hamrud, Z. Zhu, A. E. Dyas, A. N. Butterfield, A. Fyfe, E. Johnson, P. T. Conduit, γ-TuRC Heterogeneity Revealed by Analysis of Mozart1. Curr. Biol. 28, 2314–2323.e6 (2018).

54. H. Masuda, R. Mori, M. Yukawa, T. Toda, Fission yeast MOZART1/Mzt1 is an essential γ-tubulin complex component required for complex recruitment to the microtubule organizing center, but not its assembly. Mol. Biol. Cell 24, 2894–2906 (2013).

55. A. F. Brilot, A. S. Lyon, A. Zelter, S. Viswanath, A. Maxwell, M. J. MacCoss, E. G. Muller, A. Sali, T. N. Davis, D. A. Agard, CM1-driven assembly and activation of yeast γ-tubulin small complex underlies microtubule nucleation. Elife 10 (2021).

56. C. Akıl, S. Ali, L. T. Tran, J. Gaillard, W. Li, K. Hayashida, M. Hirose, T. Kato, A. Oshima, K. Fujishima, L. Blanchoin, A. Narita, R. C. Robinson, Structure and dynamics of Odinarchaeota tubulin and the implications for eukaryotic microtubule evolution. Sci. Adv. 8, eabm2225 (2022).

57. S. Hoshino, I. Hayashi, Filament formation of the FtsZ/tubulin-like protein TubZ from the Bacillus cereus pXO1 plasmid. J. Biol. Chem. 287, 32103–32112 (2012).

58. T. Matsui, J. Yamane, N. Mogi, H. Yamaguchi, H. Takemoto, M. Yao, I. Tanaka, Structural reorganization of the bacterial cell-division protein FtsZ from Staphylococcus aureus. Acta Crystallogr. D Biol. Crystallogr. 68, 1175–1188 (2012).

59. B. P. Brylawski, M. Caplow, Rate for nucleotide release from tubulin. J. Biol. Chem. 258, 760–763 (1983).

60. P. Guichard, D. Chrétien, S. Marco, A.-M. Tassin, Procentriole assembly revealed by cryo-electron tomography. EMBO J. 29, 1565–1572 (2010).

61. S. Ma, L. Li, Z. Li, S. Luo, Q. Liu, W. Du, B. Qiu, M. Gui, X. Zhu, Q. Guo, In situ cryo-electron tomography reveals the progressive biogenesis of basal bodies and cilia in mouse ependymal cells. Nat. Commun. 16, 5932 (2025).

62. B. Cai, J. Xu, E. H. Collet, E. Aarts, L. Luo, A. Leitner, T. Ishikawa, P. Beltrao, C. G. Pearson, M. Pilhofer, M. Wieczorek, Structure and assembly of the A-C linker connecting microtubule triplets in centrioles. Sci. Adv. 11, eady3689 (2025).

63. G. A. Dokshin, K. S. Ghanta, K. M. Piscopo, C. C. Mello, Robust genome editing with short single-stranded and long, partially single-stranded DNA donors in Caenorhabditis elegans. Genetics 210, 781–787 (2018).

64. M. Eroglu, B. Yu, W. B. Derry, Efficient CRISPR/Cas9 mediated large insertions using long single-stranded oligonucleotide donors in C. elegans. FEBS J. 290, 4429–4439 (2023).

65. S. Brenner, The genetics of Caenorhabditis elegans. Genetics 77, 71–94 (1974).

66. M. Ohta, O. Arakawa, Y. Gu, W. Tian, K. D. Corbett, A. Desai, K. Oegema, Phosphorylation remodels the mitotic centrosome matrix to generate bipartite γ-tubulin complex docking sites. Sci. Adv. 12, eaed6539 (2026).

67. J. B. Woodruff, O. Wueseke, V. Viscardi, J. Mahamid, S. D. Ochoa, J. Bunkenborg, P. O. Widlund, A. Pozniakovsky, E. Zanin, S. Bahmanyar, A. Zinke, S. H. Hong, M. Decker, W. Baumeister, J. S. Andersen, K. Oegema, A. A. Hyman, Centrosomes. Regulated assembly of a supramolecular centrosome scaffold in vitro. Science 348, 808–812 (2015).

68. C. Bieniossek, T. J. Richmond, I. Berger, MultiBac: multigene baculovirus-based eukaryotic protein complex production. Curr. Protoc. Protein Sci. 51, 5–20 (2008).

69. J. Cox, M. Mann, MaxQuant enables high peptide identification rates, individualized p.p.b.-range mass accuracies and proteome-wide protein quantification. Nat. Biotechnol. 26, 1367–1372 (2008).

70. A. Punjani, J. L. Rubinstein, D. J. Fleet, M. A. Brubaker, cryoSPARC: algorithms for rapid unsupervised cryo-EM structure determination. Nat. Methods 14, 290–296 (2017).

71. T. Bepler, K. Kelley, A. J. Noble, B. Berger, Topaz-Denoise: general deep denoising models for cryoEM and cryoET. Nat. Commun. 11, 5208 (2020).

72. P. Emsley, B. Lohkamp, W. G. Scott, K. Cowtan, Features and development of Coot. Acta Crystallogr. D Biol. Crystallogr. 66, 486–501 (2010).

73. P. V. Afonine, B. K. Poon, R. J. Read, O. V. Sobolev, T. C. Terwilliger, A. Urzhumtsev, P. D. Adams, Real-space refinement in PHENIX for cryo-EM and crystallography. Acta Crystallogr D Struct Biol 74, 531–544 (2018).

74. E. F. Pettersen, T. D. Goddard, C. C. Huang, E. C. Meng, G. S. Couch, T. I. Croll, J. H. Morris, T. E. Ferrin, UCSF ChimeraX: Structure visualization for researchers, educators, and developers. Protein Sci. 30, 70–82 (2021).

75. D. Kimanius, K. Jamali, M. E. Wilkinson, S. Lövestam, V. Velazhahan, T. Nakane, S. H. W. Scheres, Data-driven regularization lowers the size barrier of cryo-EM structure determination. Nat. Methods 21, 1216–1221 (2024).

76. A. Burt, B. Toader, R. Warshamanage, A. von Kügelgen, E. Pyle, J. Zivanov, D. Kimanius, T. A. M. Bharat, S. H. W. Scheres, An image processing pipeline for electron cryo-tomography in RELION-5, bioRxiv (2024)p. 2024.04.26.591129.

77. A. Hyman, D. Drechsel, D. Kellogg, S. Salser, K. Sawin, P. Steffen, L. Wordeman, T. Mitchison, “[39] Preparation of modified tubulins” in Methods in Enzymology (Elsevier, 1991)vol. 196, pp. 478–485.

78. M. Castoldi, A. V. Popov, Purification of brain tubulin through two cycles of polymerization-depolymerization in a high-molarity buffer. Protein Expr. Purif. 32, 83–88 (2003).

79. J. Schindelin, I. Arganda-Carreras, E. Frise, V. Kaynig, M. Longair, T. Pietzsch, S. Preibisch, C. Rueden, S. Saalfeld, B. Schmid, J.-Y. Tinevez, D. J. White, V. Hartenstein, K. Eliceiri, P. Tomancak, A. Cardona, Fiji: an open-source platform for biological-image analysis. Nat. Methods 9, 676–682 (2012).

80. A. Stamatakis, RAxML version 8: a tool for phylogenetic analysis and post-analysis of large phylogenies. Bioinformatics 30, 1312–1313 (2014).

81. F. D. Ciccarelli, T. Doerks, C. von Mering, C. J. Creevey, B. Snel, P. Bork, Toward automatic reconstruction of a highly resolved tree of life. Science 311, 1283–1287 (2006).

82. A. M. Altenhoff, A. Warwick Vesztrocy, C. Bernard, C.-M. Train, A. Nicheperovich, S. Prieto Baños, I. Julca, D. Moi, Y. Nevers, S. Majidian, C. Dessimoz, N. M. Glover, OMA orthology in 2024: improved prokaryote coverage, ancestral and extant GO enrichment, a revamped synteny viewer and more in the OMA Ecosystem. Nucleic Acids Res. 52, D513–D521 (2024).

83. F. Tegenfeldt, D. Kuznetsov, M. Manni, M. Berkeley, E. M. Zdobnov, E. V. Kriventseva, OrthoDB and BUSCO update: annotation of orthologs with wider sampling of genomes. Nucleic Acids Res. 53, D516–D522 (2025).

84. A. Rajković, M. Beracochea, A. B. Rogers, S. R. Eddy, N. P. Carter, R. D. Finn, HMMER web server: 2026 update. Nucleic Acids Res. 54, W272–W278 (2026).

85. P. J. A. Cock, T. Antao, J. T. Chang, B. A. Chapman, C. J. Cox, A. Dalke, I. Friedberg, T. Hamelryck, F. Kauff, B. Wilczynski, M. J. L. de Hoon, Biopython: freely available Python tools for computational molecular biology and bioinformatics. Bioinformatics 25, 1422–1423 (2009).

86. W. Kabsch, C. Sander, Dictionary of protein secondary structure: pattern recognition of hydrogen-bonded and geometrical features. Biopolymers 22, 2577–2637 (1983).

87. K. Katoh, D. M. Standley, MAFFT multiple sequence alignment software version 7: improvements in performance and usability. Mol. Biol. Evol. 30, 772–780 (2013).

88. UniProt Consortium, UniProt: The universal protein knowledgebase in 2023. Nucleic Acids Res. 51, D523–D531 (2023).

